# GIV/Girdin and Exo70 Constitute the Core of the Mammalian Polarized Exocytic Machinery

**DOI:** 10.1101/870071

**Authors:** Cristina Rohena, Navin Rajapakse, I-Chung Lo, Peter Novick, Debashis Sahoo, Pradipta Ghosh

## Abstract

Polarized exocytosis is a fundamental process by which membrane and cargo proteins are delivered to the plasma membrane with precise spatial control; it is essential for cell growth, morphogenesis, and migration. Although the need for the octameric exocyst complex is conserved from yeast to humans, what imparts spatial control is known only in yeast, i.e., a polarity scaffold without mammalian homolog, called Bem1p. We demonstrate that polarity scaffold GIV/Girdin fulfills the key criteria and functions of its yeast counterpart Bem1p. Both Bem1p and GIV bind yeast and mammalian Exo70 proteins *via* similar short-linear interaction motifs, but each preferentially binds its evolutionary counterpart. In cells where this GIV•Exo-70 interaction is selectively disrupted, delivery of the metalloprotease MT1-MMP to podosomes, collagen degradation and haptotaxis through basement membrane matrix were impaired. GIV’s interacting partners reveal other components of polarized exocytosis in mammals. Findings not only expose how GIV “upgrades” the exocytic process in mammals, but also how the ability to regulate exocytosis shapes GIV’s ability to fuel metastasis.

**GRAPHIC ABSTRACT:** **Graphic Abstract:** Schematic comparing the components of polarized exocytosis, i.e., the major polarity scaffold in yeast (Bem1p; left) and humans (Girdin; right) and the various cellular components and signaling mechanisms that are known to converge on them.

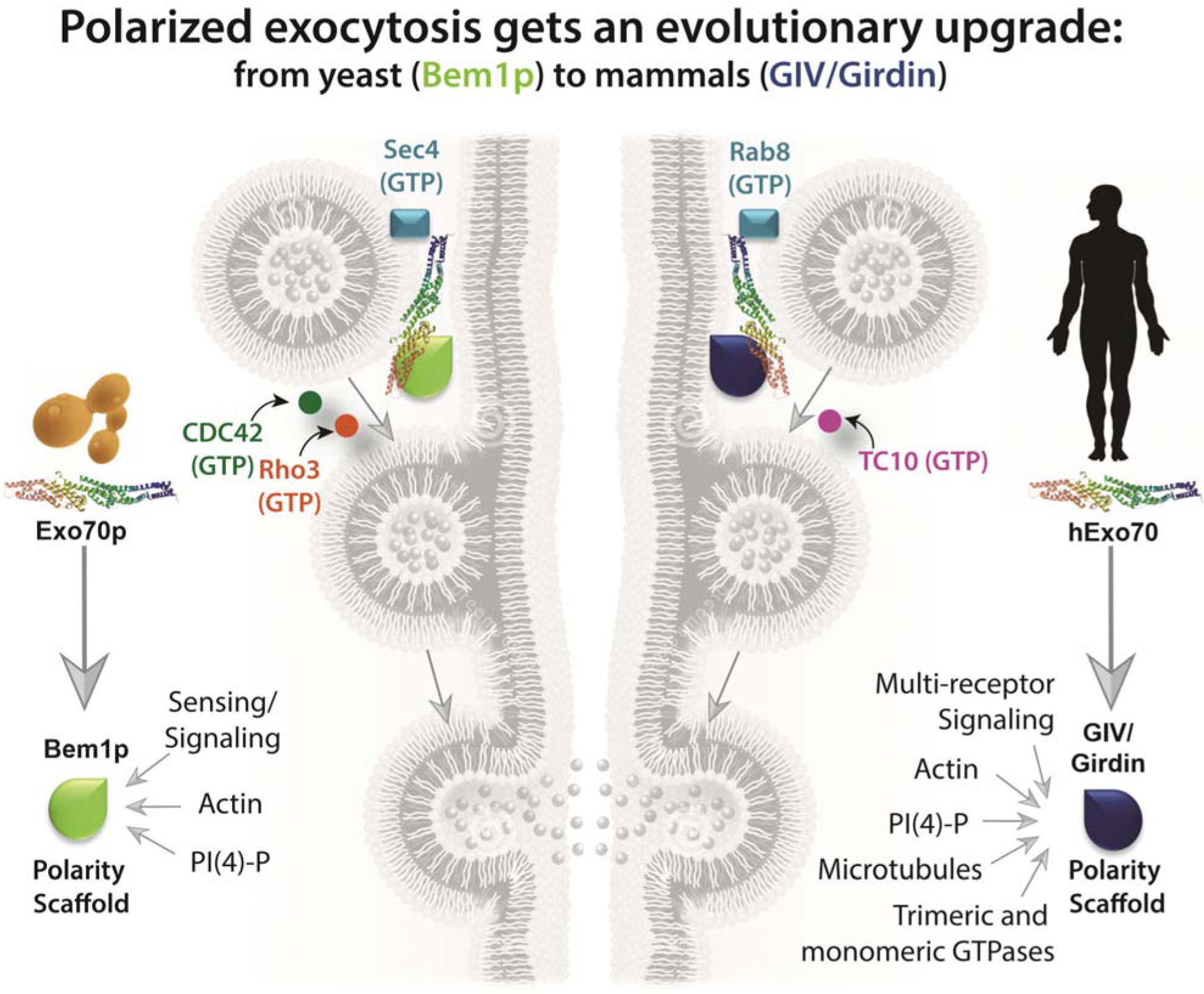

**The eTOC blurb:** Polarized exocytosis is a precision-controlled process that is enhanced in disease states, e.g., cancer invasion; what imparts polarity was unknown. Authors reveal how the process underwent an evolutionary upgrade from yeast to humans by pinpointing GIV/Girdin as the polarity scaffold which orchestrates the exocytosis of matrix metalloproteases during cell invasion.

**HIGHLIGHTS:**

- GIV (human) and Bem1p (yeast) bind Exo70; are required for exocytosis
- GIV binds and aids PM localization Exo70 *via* a conserved short linear motif
- Binding facilitates MT1-MMP delivery to podosomes, ECM degradation, invasion
- Regulatory control over polarized exocytosis is upgraded during evolution

## INTRODUCTION

Exocytosis is an essential process in all eukaryotic cells, defined as the delivery of intracellular contents such as hormones, mRNAs, and proteins between secretory vesicles and the plasma membrane (PM). The spatially controlled fusion of these secretory vesicles with the PM requires SNARE proteins. However, prior to vesicle fusion with the PM, vesicles must first be anchored to the membrane by the octameric exocyst complex. First identified in *S. cerevisiae*, using a variety of biochemical and genetic approaches (Novick et al., 1980; TerBush et al., 1996; TerBush and Novick, 1995), the exocyst complex is an evolutionarily conserved macromolecular complex comprised of Sec3, Sec5, Sec6, Sec8, Sec10, Sec15, Exo70 and Exo84. Evidence suggests that it is this exocyst complex that imparts spatial and temporal control of exocytosis, and that such precision-control is critical for a myriad of cellular processes, such as morphogenesis, cell cycle progression, primary ciliogenesis, cell migration and tumor invasion (Wu and Guo, 2015).

When it comes to spatial control, it appears that the process of exocytosis is highly polarized. Yeast genetics, biochemical (Boyd et al., 2004) and structural studies using cryo-EM and 3D molecular modeling (Mei et al., 2018) in yeast have suggested that the exocyst complex is assembled in a hierarchical manner, and that several such complexes may cooperatively ensure the fidelity in recruiting the vesicle to the appropriate exocytic site on the PM (Picco et al., 2017). For example, in the budding yeast, the exocyst is localized to the emerging bud tip, where it mediates exocytosis for the asymmetric expansion of daughter cell surfaces during polarized cell growth. During cytokinesis, the exocyst is localized to the mother–daughter cell junction to mediate abscission (Finger et al., 1998; Guo et al., 1999; TerBush and Novick, 1995; Zhang et al., 2008). Exo70p and Sec3p were identified as the key subunits of the octameric complex which plays a pivotal role in targeting of the exocyst to specific sites on the PM (Boyd et al., 2004). Such polarized localization of Exo70p and Sec3p cannot be attributed to t-SNARE (target-Soluble NSF-Attachment Protein Receptor) proteins, because the latter distributed along the entire PM (Brennwald et al., 1994). Who/what imparts polarity to the process of exocytosis remained unknown until systematic studies employing Exo70p mutants and yeast genetics pinpointed that a specific stretch of Exo70p (i.e., domain C) must interact with some other actin-independent polarity determinant (Hutagalung et al., 2009), which was subsequently identified as the multidomain polarity scaffold and bud emergence protein, Bem1p (Liu and Novick, 2014). This Exo70p•Bem1p interaction was implicated in polarizing the process of vesicle exocytosis in yeast; binding-deficient mutants of Exo70p were incapable of tethering secretory vesicles to cortical sites specified by Bem1p (Liu and Novick, 2014).

Although the fundamentals of exocytosis were discovered first in yeast, its existence and importance in mammals is widely accepted as a driver in multiple disease states in humans (Martin-Urdiroz et al., 2016), insights into how the process is regulated in mammals remains unknown to date. As in yeast, the mammalian Exo70 has been the most-studied subunit; these studies have revealed that although the composition of the canonical octameric exocyst component is essentially invariant (Boehm and Field, 2019), there is however a remarkable degree of evolutionary flexibility to allow for specialization through diversity in exocytic mechanism and building more regulatory control along the way. For example, unlike yeast Exo70p, mammalian Exo70 interacts directly with Arp2/3 complex during cell migration and enhances actin cytoskeleton remodeling (Zuo et al., 2006). Mammalian Exo70 is also under the control of growth factors—EGF stimulation triggers phosphorylation of Exo70 by ERK1/2 leading to enhanced octameric complex formation, MMP secretion and cell invasion (He et al., 2007; Hertzog et al., 2012; Liu et al., 2009; Lu et al., 2013; Zhao et al., 2013). Additionally, a switch between Exo70 isoforms has been reported in breast cancer that correlates with a switch from epithelial to mesenchymal transition (EMT)(Lu et al., 2013); the former is engaged in exocytosis, whereas the latter modulates actin dynamics. The fact that Exo70 was the only exocyst subunit found to correctly localize when overexpressed in mammalian epithelial cells (He et al., 2007; Yeaman et al., 2004) suggested that Exo70 might be the critical component that recognizes a polarity determinant at the cell cortex. In the absence of any clear mammalian counterpart of yeast polarity scaffold Bem1p, it has been speculated that other mammalian polarity determinants may fulfil that role to facilitate Exo70 recruitment in mammals. The identity of such determinant remains unknown.

Here we reveal that the multi-modular polarity scaffold Gα-interacting vesicle associated protein (GIV, a.k.a, Girdin) (Aznar et al., 2016b; Ohara et al., 2012; Sasaki et al., 2015) fulfils all criteria to serve as the mammalian functional homolog of the yeast polarity determinant, Bem1p. The scaffolding function of GIV has been shown to mediate the purposeful dynamic ‘glue’-ing of diverse signaling pathways: GIV is a guanine nucleotide exchange factor (GEF) for trimeric GTPases, Gi (Garcia-Marcos et al., 2009; Kalogriopoulos et al., 2019) and Gs (Gupta et al., 2016), a polarity scaffold that binds and modulates via G protein intermediates the aPKC/Par complexes (Ohara et al., 2012; Sasaki et al., 2015); it is a substrate of and major signaling conduit for multiple receptor and non-receptor tyrosine kinases (Lin et al., 2011; Midde et al., 2018; Mittal et al., 2011), serves as a signaling adaptor for growth factor (Lin et al., 2014), integrins (Leyme et al., 2015; Lopez-Sanchez et al., 2015a) and other classes of receptors [reviewed in (Ghosh et al., 2017)], is an actin remodeler (Enomoto et al., 2005) and a scaffold for polymerized microtubules (Aznar et al., 2016b; Nechipurenko et al., 2016). Most of the work on GIV thus far have revealed its ability to serve as an integrator of multiple signaling pathways that enhances multiple fundamental cellular processes, and how its dysregulation (overexpression and hyperactivation) favors cancer invasion and metastasis [reviewed in (Aznar et al., 2016a)]. This work not just sheds light into GIV’s ability to enhance a key process that aids metastasis (i.e., polarized exocytosis of matrix metalloproteases), but also reveals how the modular makeup and functional diversity of GIV imparts the process of exocytosis features of polarity and context-dependent regulation by diverse signaling pathways.

## RESULTS AND DISCUSSION

### GIV and Bem1p share key properties, conserved modes of binding to Exo70

We first began by looking for polarity scaffold proteins with the mammalian Exo70 interactome (**Fig S1**) using Biological General Repository for Interaction Datasets (BioGRID; thebiogrid.org). A systematic analysis of all physical interactions of Exo70 revealed that among many coiled coil multi-modular proteins, it binds the mammalian polarity scaffold GIV (encoded by the gene CCDC88A) as revealed initially in a yeast 2-hybrid screen (Camargo et al., 2007) and validated later by us as a protein that colocalizes with (as determined by confocal immunofluorescence and *in situ* proximity ligation assays) and directly binds Exo70 (Lopez-Sanchez et al., 2015c). We noted that although the modular makeup of GIV and Bem1p had very little in common (**Fig 1A**), GIV fulfilled all functional criteria that were previously deemed as necessary for Bem1p to polarize exocytosis in yeast (**Fig 1B**). Furthermore, besides its ability to bind and modulate Par-polarity complexes, GIV presented numerous additional modules, functional features and interesting interactions (**Fig 1A-B**). We hypothesized that GIV may be the long-sought evolutionary counterpart of Bem1p. Because Exo70 was proposed as the exocyst subunit that interacts with polarity determinants in mammalian cells (He et al., 2007; Yeaman et al., 2004), we began first by comparing the abilities of Bem1p and GIV to bind yeast and human Exo70. We found that both GST-tagged human (hExo70) and yeast (Exo70p) Exo70 could bind full length endogenous GIV (**Fig S2A**). Using recombinant His-tagged proteins in similar pulldown assays with GST-Exo70, we confirmed that the GIV•Exo70 interaction is direct and that the C-terminus of GIV is sufficient (GIV-CT; **Fig 1C**). While both Bem1p and GIV-CT could bind both yeast and human Exo70 proteins, each preferentially bound to its evolutionary counterpart (**Fig 1C-D**): GIV-CT binds four-fold as strongly to human Exo70 compared to yeast Exo70p (**Fig 1C**), whereas Bem1p binds two-fold as strongly to yeast Exo70p compared to human Exo70 (**Fig 1D**).

**Figure 1.**
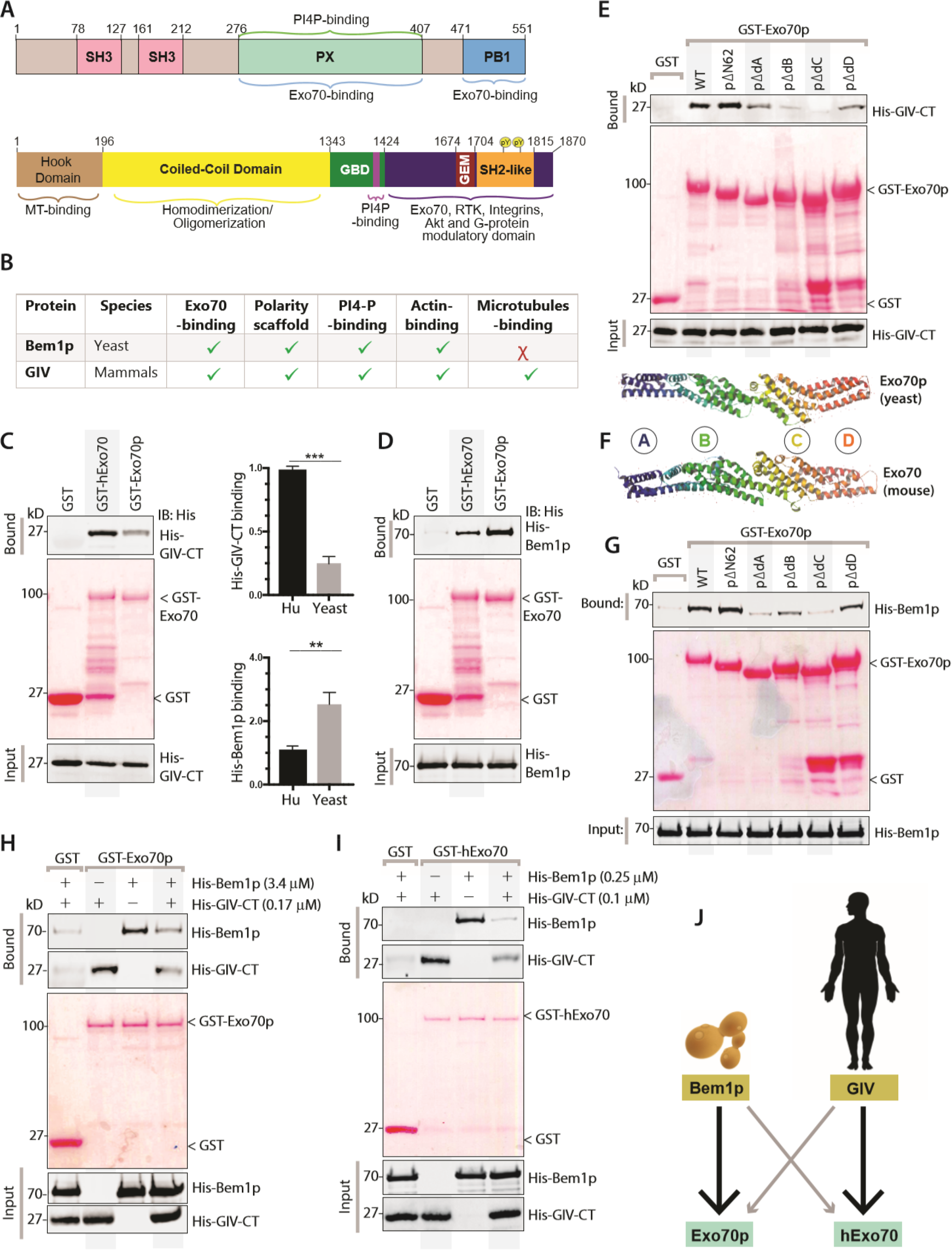
Polarity scaffolds GIV and Bem1p share key properties, display conserved modes of binding to Exo70. **A)** Domain maps of the yeast [Bem1p (top)] and human [GIV/Girdin (bottom)] polarity scaffolds. Major interactions are annotated. **B)** Table listing key properties of Bem1p that have been implicated in its ability to impart polarity to the process of exocytosis. **C)** Recombinant His-GIV-CT (~ 3 μg) was used in GST pulldown assays with GST, GST-hExo70 (human) and GST-Exo70p (yeast). Bound GIV was visualized by immunoblotting using anti-His mAb. Equal loading of GST proteins was confirmed by Ponceau S staining. *<u>Top right</u>*: Bar graph displays fold change in binding. N = 3; *** = 0.001. **D)** Recombinant His-Bem1p (~ 3 μg) was used in GST pulldown assays with GST proteins as in C. Bound Bem1p was visualized by immunoblotting using anti-His mAb. Equal loading of GST proteins was confirmed by Ponceau S staining. *<u>Bottom left</u>*: Bar graph displays fold change in binding. N = 3; ** = 0.01. **E-G)** Pulldown assays as in C were carried out using either recombinant His-GIV-CT (**E**) or His-Bem1p (**G**) and various GST constructs (Liu and Novick, 2014) lacking the indicated domains of Exo70 (yeast vs. mouse comparison; **F**) immobilized on glutathione beads. Bound GIV (**E**) or Bem1p (**G**) were visualized by immunoblotting using anti-His mAb. Equal loading of GST proteins was confirmed by Ponceau S staining. **H-I)** Competition between GIV and Bem1p for binding to either Exo70p (yeast; H) or hExo70 (human; I) was assessed by carrying out pulldown assays adding (+) or not (-) the soluble His-GIV-CT and His-Bem1p proteins alone or simultaneously to GST-Exo70 proteins immobilized on glutathione beads. Bound GIV or Bem1p proteins were visualized by immunoblotting using anti-His mAb. **I)** Schematic summarizing the cross-species binding of Bem1p and GIV to Exo70. The size and darkness of arrows indicate binding preference.

Prior work showed that although Bem1p forms an extended interface with Exo70p, domain C is the most critical part of Exo70 that is responsible for the interaction (Liu and Novick, 2014). Using a series of previously validated Exo70p mutants, we confirmed that both Bem1p and GIV-CT form an extensive and similar interface with Exo70—although domain C is the major contributor, domains A, B and D are also relevant (**Fig 1E-G**). Despite the overall similarity in domain contributions, combination of single point mutations on Exo70p that disrupted the Bem1p•Exo70 interaction (Liu and Novick, 2014) did not affect binding to GIV-CT (**Fig S2B**). These findings are in keeping with the fact that *mammalian and yeast Exo70* differs in both conformation and surface charge (**Fig 1F, S2C**) and share low level of sequence conservation which has been suggested as a basis for species-specific functions of the exocyst (Moore et al., 2007). In the absence of conserved amino acids across species, or any other structural clues, generating corresponding mutations in human Exo70 was not attempted. When we carried out competition assays with recombinant proteins to determine if the Bem1p•Exo70p and GIV•Exo70 interfaces are similar. We found that both GIV and Bem1p could compete for yeast (**Fig 1H**) and human (**Fig 1I**) Exo70 demonstrating that they not only share the mechanism of binding, but also that their binding interfaces with Exo70 are significantly overlapping. Taken together, these findings demonstrate that despite divergent modular makeup, both polarity scaffolds Bem1p (yeast) and GIV (mammals) can bind Exo70 via mechanisms that are shared, using interfaces that are similar; although each can bind both yeast and human Exo70 proteins, they demonstrate species-preference (**Fig 1J**).

### GIV is required for Exo70 localization, polarized exocytosis

Because GIV’s role in tumor cell invasion and its ability to promote cancer metastasis has been demonstrated *in vitro* (Jiang et al., 2008; Leyme et al., 2015; Rahman-Zaman et al., 2018) and *in vivo* (Jiang et al., 2008) using MDA MB231 human breast cancer cells, we used the same cells and asked if GIV is required for the polarized delivery of cargo proteins e.g., matrix metalloproteases that aid in tumor dissemination by degrading extracellular matrix (ECM). We first confirmed that depletion of GIV (~85% depletion by shRNA; **Fig 2B**) does not change the levels of Exo70 or cortactin; the latter is a marker of podosomes, which are subcellular actin-rich landmarks that are specialized for matrix degradation because of polarized exocytosis of proteinases (Castro-Castro et al., 2016; Hoshino et al., 2013). We found that HA-tagged Exo70 localized to the PM in control but not GIV-depleted cells (**Fig 2B, S3A**, top panel). In control cells, disruption of actin cytoskeleton with latrunculin B, a chemical that has been widely used for its ability to sequester G-actin and prevent F-actin assembly (Morton et al., 2000) prevented fusion of Exo70-positive vesicles but did not impact their docking at specific sites on the PM; in GIV-depleted cells the vesicles were in disarray (**Fig 2B, S3A**, bottom panel; **Fig 2C**). These findings indicate that GIV is required for the localization of Exo70 to the PM, and that such localization is actin independent.

**Figure 2.**
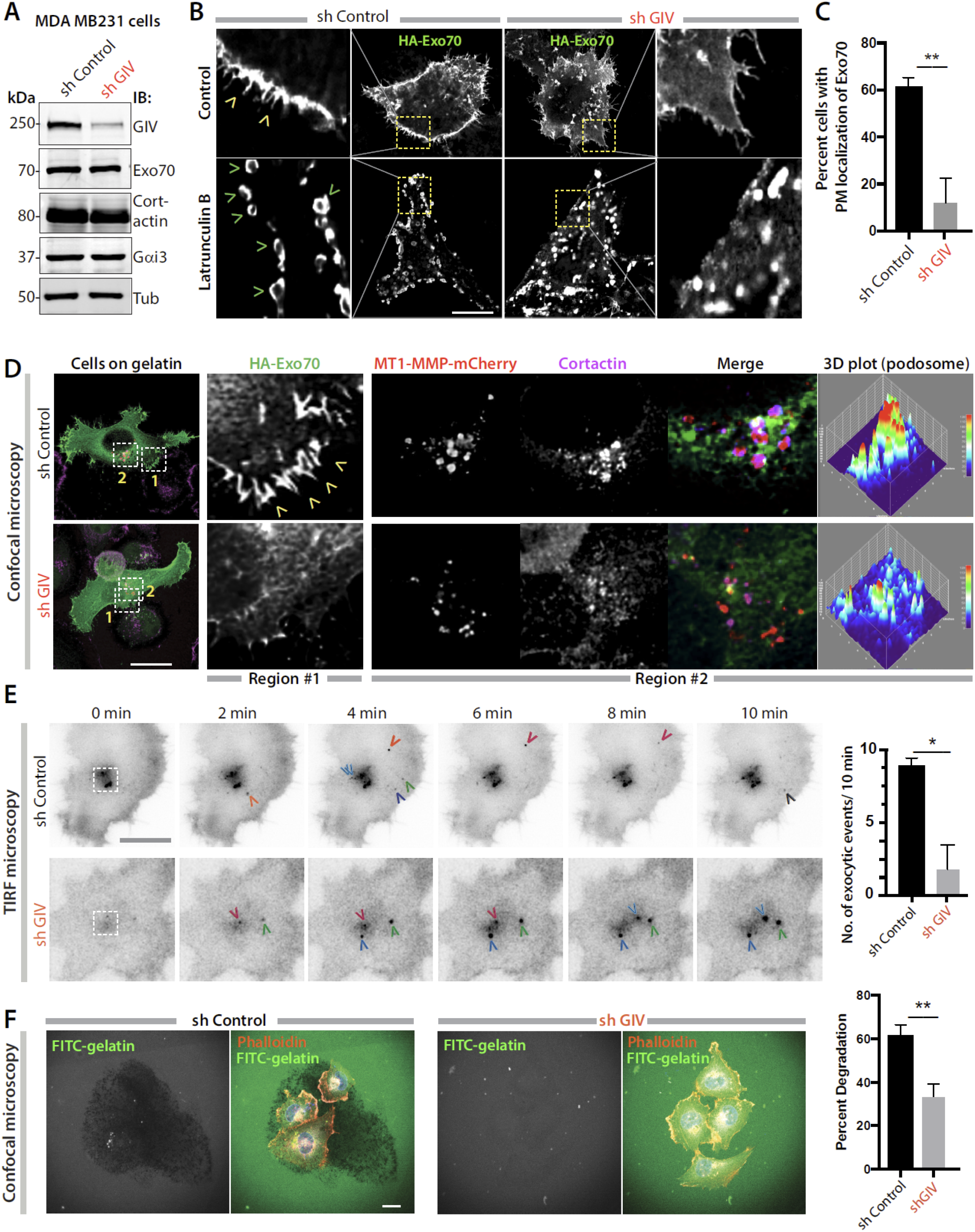
GIV is required for Exo70 localization, polarized exocytosis. **A)** Immunoblots of lysates of control (sh Control) and GIV-depleted (sh GIV) MDA MB231 cells. **B)** control (sh Control) and GIV-depleted (sh GIV) MDA MB231 cells expressing HA-Exo70 were treated (bottom) or not (top) with 25 μM Latrunculin B for 24 h prior to fixation with PFA. Fixed cells were stained for HA and analyzed by confocal microscopy. Representative deconvolved images are shown. Arrowheads = PM localization. **C)** Bar graphs show the % cells with PM-localization of Exo70-HA. N = ~ 30 cells/assay. Error bars = S.E.M. *p* values, ** < 0.01. **D)** Cells in B exogenously expressing HA-Exo70 (green) and MT1-MMP-mCherry (red) were plated on gelatin-coated coverslips, fixed and stained with the podosome marker cortactin (magenta). Representative deconvolved images are shown. Insets were analyzed by rendering 3D surface plots using ImageJ. **E)** Cells in B exogenously expressing MT1-MMP-pHLuorin were plated on a layer of unlabeled gelatin and imaged by TIRF microscopy. *<u>Left</u>*: Still frames from a 10 min long movie (*Supplementary movies 1, 2*). Each colored arrowhead marks one successfully exocytosed vesicle. *<u>Right</u>*: Bar graph shows the number of exocytic events that were encountered during the 10-min long movie in D. N = ~ 5-10 cells/assay. Error bars = S.E.M. *p* values, * < 0.05. **F)** Cells in B were plated for 5 h on FITC-conjugated cross-linked gelatin (green), and then fixed and stained for F-actin (Phalloidin; red). *<u>Left</u>*: Representative images are shown. *<u>Right</u>*: Bar graphs display the % of cells that showed degradation of gelatin. N = ~100-200 cells /experiment x 3. Error bars = S.E.M. *p* values, ***0.001.

Next, to analyze how Exo70 localization may impact polarized exocytosis of proteases to podosomes, we co-transfected the MDA MB231 cells with HA-Exo70 and a previously validated MT1-MMP-mCherry construct (Steffen et al., 2008), plated them on gelatin and looked for the co-localization of MT1-MMP, Exo70 and cortactin. Again, HA-Exo70 localized to the PM in the presence of GIV, but significantly less in its absence (**Fig 1D**; Region #1). Colocalization between MT1-MMP-mCherry and cortactin was easily detected in control cells, but significantly reduced in cells without GIV (**Fig 1D, S3B**; Region #2 and 3D plot of podosome). These findings indicate that GIV is required for efficient localization of proteases near podosomes. To assess the dynamics of polarized exocytosis in real time, we used a pH-sensitive MT1-MMP-pHluorin construct (Monteiro et al., 2013), which is fluorescent only at an extracellular pH of 7.4. Prior studies using this construct and total internal reflection fluorescence (TIRF) live-cell microscopy have demonstrated that MT1-MMP-pHluorin demonstrates dot-like localization at the substrate-attached cell side, reminiscent of podosomes (El Azzouzi et al., 2016; Planchon et al., 2018). When we carried out TIRF-microscopy on control and GIV-depleted MDA MB231 cells, we found that the number of exocytic events were significantly lower in the latter (**Fig 1E**), and that they were not clustered near the center of the cell (**Fig 1E**; interrupted white square). Consistent with these findings, compared to controls cells fewer % of GIV-depleted cells demonstrated gelatin degradation (**Fig 1F**). Together, these findings suggest that GIV is required for the localization of Exo70 to specific sites at the PM, and for the polarized exocytosis of MT1-MMP at the podosomes. Findings also suggest that GIV-dependent exocytosis of the protease may be important for the efficient degradation of ECM.

### A short linear motif on C-terminus mediates GIV’s interaction with Exo70

To pinpoint the role of GIV in polarized exocytosis, we set out to identify mutant(s) of GIV that may not bind Exo70. Previously, using domain mapping studies the Exo70p-binding domain of Bem1p was narrowed down to amino acids 309–510, which includes three quarters of the PX domain, half of the PB1 domain, and the region connecting them (see **Fig 1A**) as both necessary and sufficient (Liu and Novick, 2014). Because Bem1p and GIV appeared to share the mechanism of binding to Exo70, via what appears to be extensive binding interfaces (**Fig 1H-I**), we hypothesized that the interactions could be sensitive to some key residues. When aligned Bem1p and GIV, we noticed that they have very little similarity in sequences, except for a short stretch of sequence within the PX domain of Bem1p, ^320^DFYD^323^ (**Fig 3A-B**). When we superimposed two previously solved structures of Bem1p, we noted that the Phenylalanine (F) within this sequence exhibits flexibility (**Fig 3A**). We targeted this putative short linear interaction motif (SLIM, ^1741^DFYD^1744^) in GIV by either selectively replacing the Phe>Ala (F^1742^A) or by replacing the motif with Ala at 3 of the 4 positions (AAxA). Both mutants did not bind GST-Exo70 (**Fig 3C**). Next we carried out a series of mutations within and flanking the SLIM and noted that several impaired binding of GIV to Exo70 (**Fig 3D**), demonstrating that regardless of potentially extensive contact sites, the GIV•Exo70 interaction is sensitive to point directed mutagenesis.

**Figure 3.**
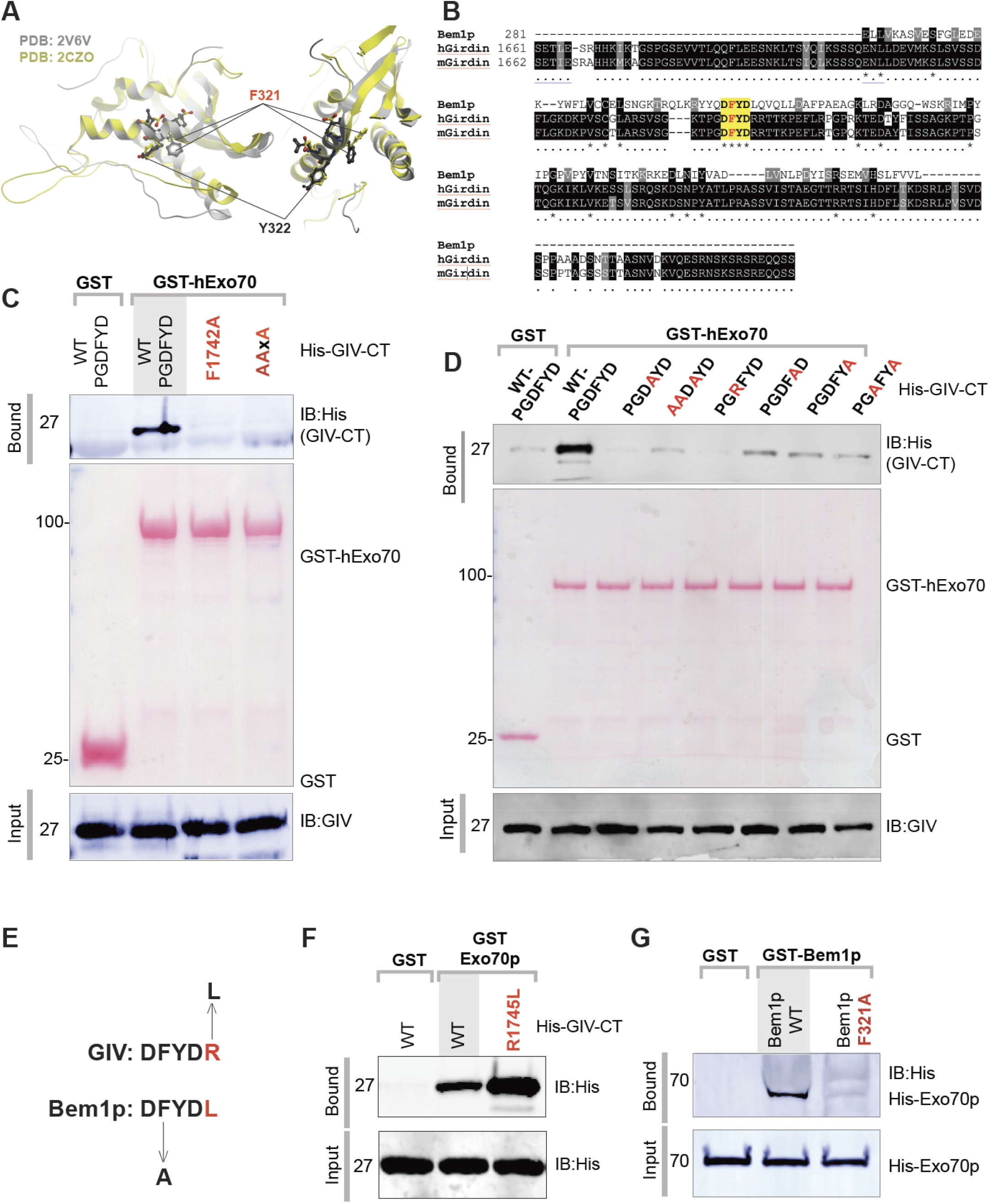
A short linear motif on C-terminus mediates GIV’s interaction with Exo70. **A)** Two views of the two solved crystal structures of PX domain of Bem1p [PDB:2V6V, (Stahelin et al., 2007); PDB:2CZO, *unpublished*], showing the position and orientation of key residues Phe (F)321 and Tyr (Y)322 on Bem1p that are within the previously identified stretch of amino acids 309–510 that is both necessary and sufficient for binding to Exo70p. **B)** Sequence alignment showing a short linear ‘DFYD’ motif that is conserved between GIV and Bem1p. **C-D)** Recombinant His-GIV-CT (~ 3 μg) was used in GST pulldown assays with GST or GST-hExo70 (human) WT and mutant proteins. Bound GIV was visualized by immunoblotting using anti-His mAb. Equal loading of GST proteins was confirmed by Ponceau S staining. **E)** Sequence-based prediction of mutations in GIV (top; R1742»L) and Bem1p (bottom; F312»A) that should enhance and reduce binding to GST-Exo70p (yeast), respectively. **F)** Recombinant His-GIV-CT WT or R1742»L mutant was used in GST pulldown assays with GST or GST-hExo70 (human). Bound GIV was visualized by immunoblotting using anti-His mAb. **G)** Recombinant His-Exo70p was used in GST pulldown assays with GST or GST-Bem1p WT or F312»A mutant. Bound Exo70p was visualized by immunoblotting using anti-His mAb.

The sequence alignment also predicted mutations that would improve GIV’s ability to bind yeast Exo70p (i.e., R^1745^L) or those that might impair Bem1p’s ability to bind Exo70p (F^321^A) (**Fig 3E**). We found both predictions to be true—a mutant GIV (R^1745^L) bound Exo70p stronger than GIV-WT (**Fig 3F, S4A**). Similarly, the F321A mutant Bem1p did not bind Exo70p (**Fig 3G**). These findings provide further evidence that while the interactions between GIV or Bem1p is complex and may occur *via* an extensive interface, and that the mechanisms of these interactions in yeast and mammals are divergent for the most part, the newly identified SLIM may represent an evolutionary conserved mechanism (**Fig S4B**). This Exo70-binding SLIM, localized within the C-terminus of GIV, adds to the gathering catalog of SLIMs used by GIV to scaffold diverse signaling components (**Fig S4C**).

### Exo70 binding-deficient GIV mutant fails to deliver MT1-MMP to podosomes

We next asked if the phenotypes we observed previously (**Fig 2**) were because of GIV’s ability to bind Exo70. To this end, we generated MDA MB231 cell lines in which endogenous GIV was depleted by shRNA before being restored with full length GIV either WT or the newly identified Exo70 binding-deficient GIV mutant AAxA (**Fig 4A**) and used them in similar assays as before. We found that compared to cells expressing GIV-WT, those expressing the binding-deficient AAxA GIV mutant showed impaired localization of HA-Exo70 at the PM (**Fig 4B**), colocalization of both Exo70 and MT1-MMP-mCherry to cortactin-positive podosomes (**Fig 4C**), rate of exocytic events, as determined by TIRF microscopy on cells expressing MT1-MMP-pHluorin (**Fig 4D-E**) and gelatin degradation (**Fig 4F-G**). These results were also validated using another Exo70-binding deficient mutant, F^1742^A (**Fig S5A-C; S6**). There findings indicate that the GIV•Exo70 interaction may be critical for the localization of Exo70 to vesicle tethering sites at the PM and for the polarized exocytosis of MT1-MMP proteases at/near the podosomes of cells.

**Figure 4.**
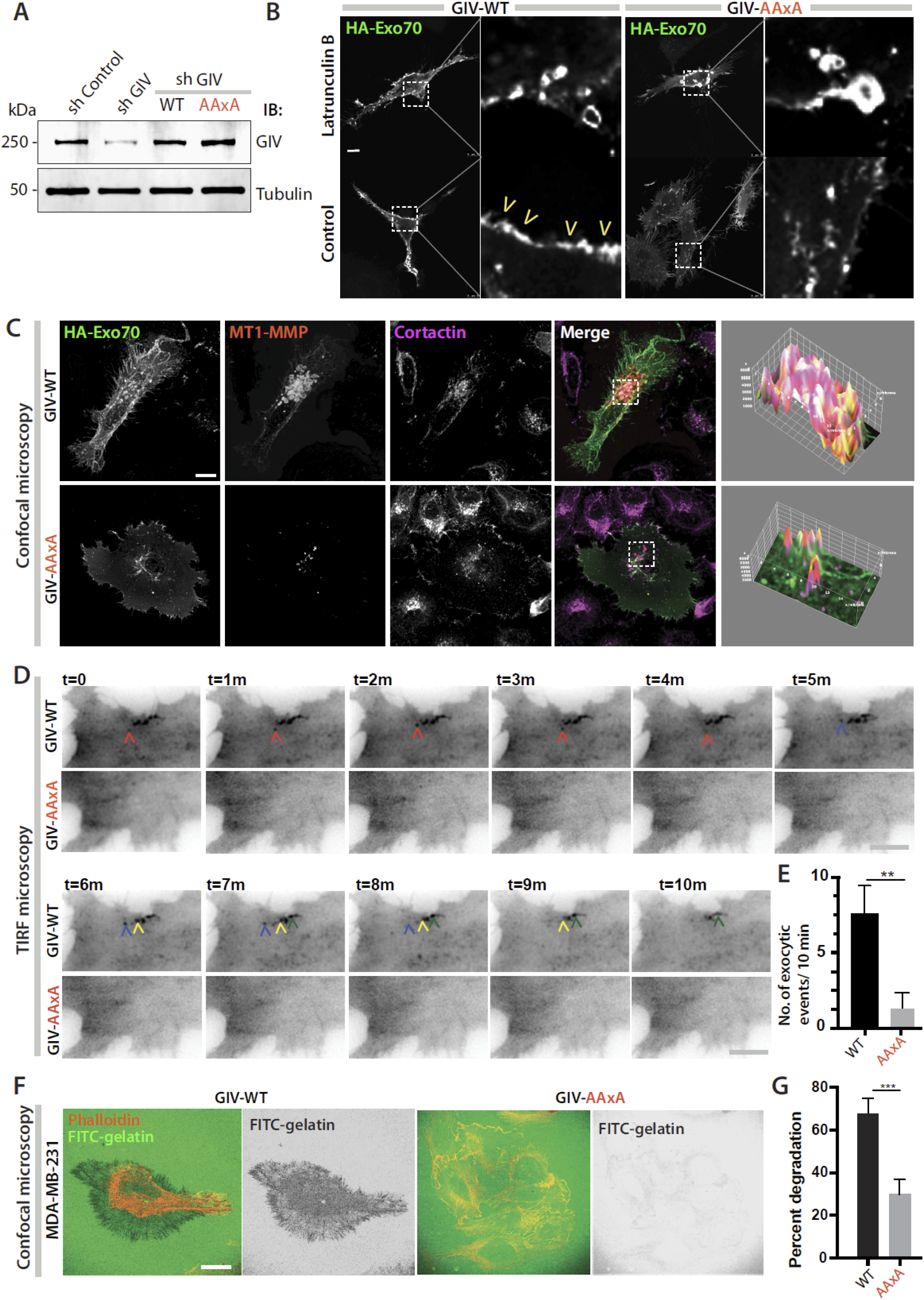
Exo70-binding deficient mutant GIV fails to deliver MT1-MMP to podosomes. **A)** Immunoblots of lysates of control (sh Control) and GIV-depleted (sh GIV) MDA MB231 cells stably expressing GIV-FLAG WT and exo70-binding deficient AAxA mutant constructs. See also **Fig S5A**. **B)** GIV-depleted (sh GIV) MDA-MB-231 cells stably expressing GIV-WT or GIV-AAxA mutant were transfected with HA-Exo70 were treated with either treated (top) or not (bottom) with 25 μM Latrunculin B for 24 h prior to fixation with PFA. Fixed cells were stained for HA and analyzed by confocal microscopy. Representative deconvolved images are shown. Arrowheads = PM localization. **C)** Cells in B transfected with HA-Exo70 (green) and MT1-MMP-mCherry (red) were plated on gelatin-coated coverslips, fixed and stained with the podosome marker cortactin (magenta). Representative deconvolved images are shown. Insets were analyzed by rendering 3D surface plots using ImageJ. See also **Fig S5B** for F1742A mutant GIV. **D)** Cells in B were transfected with MT1-MMP-pHLuorin were plated on a layer of unlabeled gelatin and imaged by TIRF microscopy. *<u>Left</u>*: Still frames from a 10 min long movie (*Supplementary movies 3, 4*). See also **Fig S5C** for F1742A mutant GIV. **E)** Bar graph shows the number of exocytic events that were encountered during the 10-min long movie in D. Each colored arrowhead tracks one vesicle. N = ~ 5-10 cells/assay. Error bars = S.E.M. *p* values, **0.01. **F)** Cells in B were plated for 5 h on FITC-conjugated cross-linked gelatin (green), and then fixed and stained for F-actin (Phalloidin; red). Representative images are shown. See also **Fig S6** for F1742A mutant GIV. **G)** Bar graphs display the % of cells that showed degradation of gelatin. N = ~100-200 cells /experiment x 3. Error bars = S.E.M. *p* values, ***0.001.

### Cells expressing the Exo70-binding deficient mutant GIV fails to spread, degrade matrix, invade

Next we asked how the GIV•Exo70 interaction impacts cellular phenotypes that have been previously attributed to exocytosis of either membrane or proteases, e.g., cell spreading and invasion through the basement membrane ECM, but not impact adhesion; the latter is primarily mediated by integrins and is independent of exocytosis (**Fig 5A**). We hypothesized that cells expressing a mutant GIV that cannot bind Exo70 may adhere normally but may not spread or invade efficiently through the ECM. Cell adhesion assays confirmed that depletion of GIV impairs adhesion on collagen-coated surface (**Fig 5B**), which is in keeping with prior work by others and us showing that GIV modulates integrin signaling (Leyme et al., 2016; Leyme et al.,2015; Lopez-Sanchez et al., 2015a). Cells expressing either GIV-WT or the binding-deficient GIV-AAxA mutant adhered to the collagen to similar extents as control cells (**Fig 5B**), indicating that the cell adhesion occurs independent of the GIV•Exo70 interaction. As for cell spreading, we analyzed spreading at/beyond ~30 min, a phase of spreading that requires membrane exocytosis and is proteolysis-dependent (**Fig 5C**). We found that although both GIV-WT and GIV-AAxA cells adhered equally well to collagen-coated surfaces (**Fig 5B**), the latter showed significant impairment in cell spreading (**Fig 5D-E**), indicating that cell-spreading requires an intact GIV•Exo70 interaction. Haptotaxis assays carried out across a 0%»10% serum gradient (**Fig 5F**) showed that depletion of GIV impairs cell invasion, however, such impairment is rescued by GIV-WT, but not the Exo70 binding-deficient GIV-AAxA mutant (**Fig 5G-H**). These findings demonstrate that the GIV•Exo70 interaction is required for key sinister properties of cancer cells that require efficient polarized exocytosis of protases that degrade the ECM and aid in tumor dissemination.

**Figure 5.**
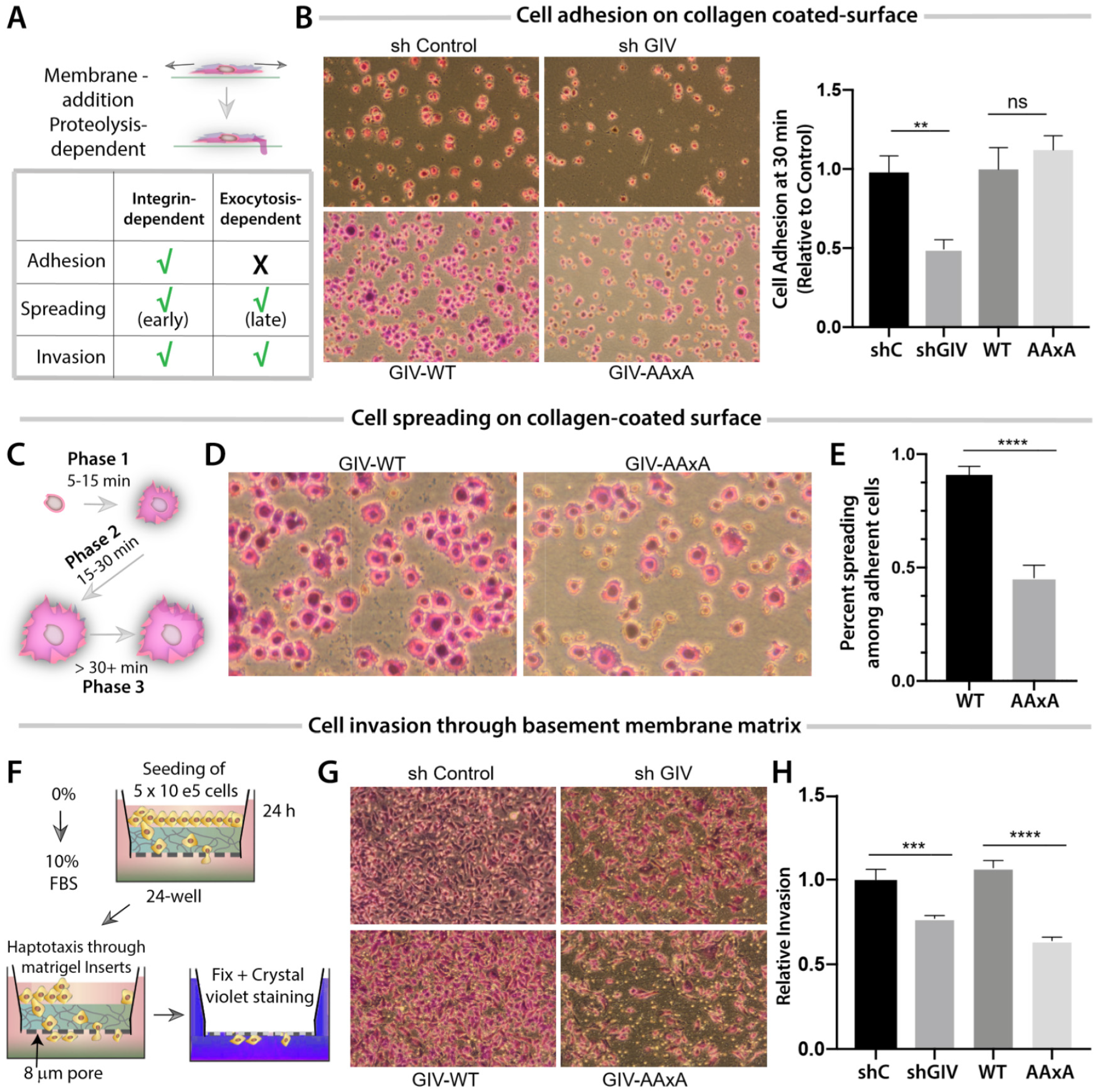
Cells expressing Exo70-binding deficient mutant GIV adhere, but fail to spread, degrade matrix, and invade. **A)** Schematic summarizing how polarized exocytosis of membrane and cargo (in this case MT1-MMP) impacts cell adhesion, spreading and invasion. **B)** MDA-MB 231 cell lines were analyzed for cell adhesion by growing on poly-D-Lysine coated surface, resuspending and then plating on 12-well collagen coated plates for 30 min before fixing with 4% PFA and staining with crystal violet. Cells were visualized and imaged by light microscopy. *<u>Left</u>*: Representative images are shown. *<u>Right</u>*: Bar graphs display the relative numbers of adherent cells per field, as determined using ImageJ. Error bar = S.E.M (n = 4); \*\**p* = < 0.01; ns = not significant. **C)** Schematic summarizing the phases of cell spreading during adhesion. **D)** Adherent MDA-MB 231 cells in B were further analyzed for attachment-induced cell-spreading at higher magnification. Representative images are shown. **E)** Bay graphs display the quantification of % spreading in D. Error bar = S.E.M (n = 4); \*\*\*\**p*=<0.0001. **F)** Schematic diagram showing the serum gradient-induced haptotaxis assay conditions used in G-H. **G)** MDA-MB 231 cell lines in A were analyzed for the ability to invade through matrigel-coated transwells. The number of cells that successfully invaded within 24 h was imaged. Representative images are shown. **H)** Bar graphs display the number of invading cells in G, as determined using Image-J. Error bar = S.E.M (n = 4); ***p=<0.001, ****p=<0.0001.

### Components of polarized exocytosis synergize during tumor metastasis

Next we asked if the observed interaction between GIV and Exo70 we observe here and the impact of such interaction on sinister properties of tumor cells can be meaningful when assessing tumor behavior and/or prognosticating clinical outcome. First we asked if the levels of expression of Exo70 or GIV alone could impact one of the most important readouts of cancer aggressiveness, i.e., metastasis-free patient survival. To discern this, we chose to study a pooled cohort of patients (Bos et al., 2009; Minn et al., 2005; Wang et al., 2005) with breast cancers who did not receive adjuvant chemotherapy, and hence, in them metastatic progression reflects natural disease progression and not resistance/selection under treatment. Samples were divided into “low” and “high” subgroups with regard to Exo70 (EXOC7) and GIV (CCDC88A) gene expression levels using the StepMiner algorithm (Sahoo et al., 2007), implemented within the hierarchical exploration of gene-expression microarrays online (HEGEMON) software (Dalerba et al., 2011; Volkmer et al., 2012). Kaplan-Meier analyses of the disease free-survival showed that Exo70 and GIV had opposing impact on outcome (**Fig 6A**)--Consistent with numerous prior studies [reviewed in (Ghosh, 2015)], high levels of GIV expression carried a grave prognosis, i.e., a shorter metastasis-free survival. By contrast, high expression of Exo70 carried a better prognosis, i.e., a longer metastasis-free survival (**Fig 6A**); this is also consistent with prior findings that an epithelial isoform of Exo70, which is the isoform that is primarily responsible for exocytic secretions, inhibits tumor metastasis in mice (Lu et al., 2013). We sought to determine if the levels of expression of either GIV or Exo70 and other genes that encode major polarity scaffolds (Lin et al., 2015) interact synergistically as independent variables to impact survival of patients. To this end, we carried out a Cox proportional-hazards model (Cox, 1972) which is a regression model that is commonly used as a statistical method for investigating the effect of several variables upon the time it takes for a specified event to happen, in this case, metastasis. We found that besides GIV/CCDC88A, Exo70 significantly and negatively interacts with multiple members of the Par, Scribble and Crumbs family of polarity scaffolds (**Fig 6B**), in that, high levels of expression of each of these scaffolds in the setting of high Exo70 expression was sufficient in shortening metastasis-free survival. Of all those polarity scaffolds that interacted with Exo70, only DLG5 continued to show positive synergistic interactions with CCDC88A (GIV) (**Fig 6C**), in that, high levels of expression of DLG5 in the setting of high GIV expression further shortened metastasis-free survival. Next we asked how Exo70 or GIV interacts with other components of the exocytic machinery, i.e., core trafficking components (i.e., monomeric GTPases Rabs, Arf6), focal adhesions cytoskeleton and Rho GTPases (Paxillin, PXN; Actin, ACTB; TC10, RHOQ; Rho-associated kinase, ROCK1), cargo (MT1-MMP, MMP14), and receptors (ITGB1 and EGFR). We found that while many components may negatively interact with Exo70 (**Fig 6D**), only TC10 GTPase (RHOQ) positively and synergistically interacts with GIV to shorten metastasis-free survival (**Fig 6E**).

**Figure 6.**
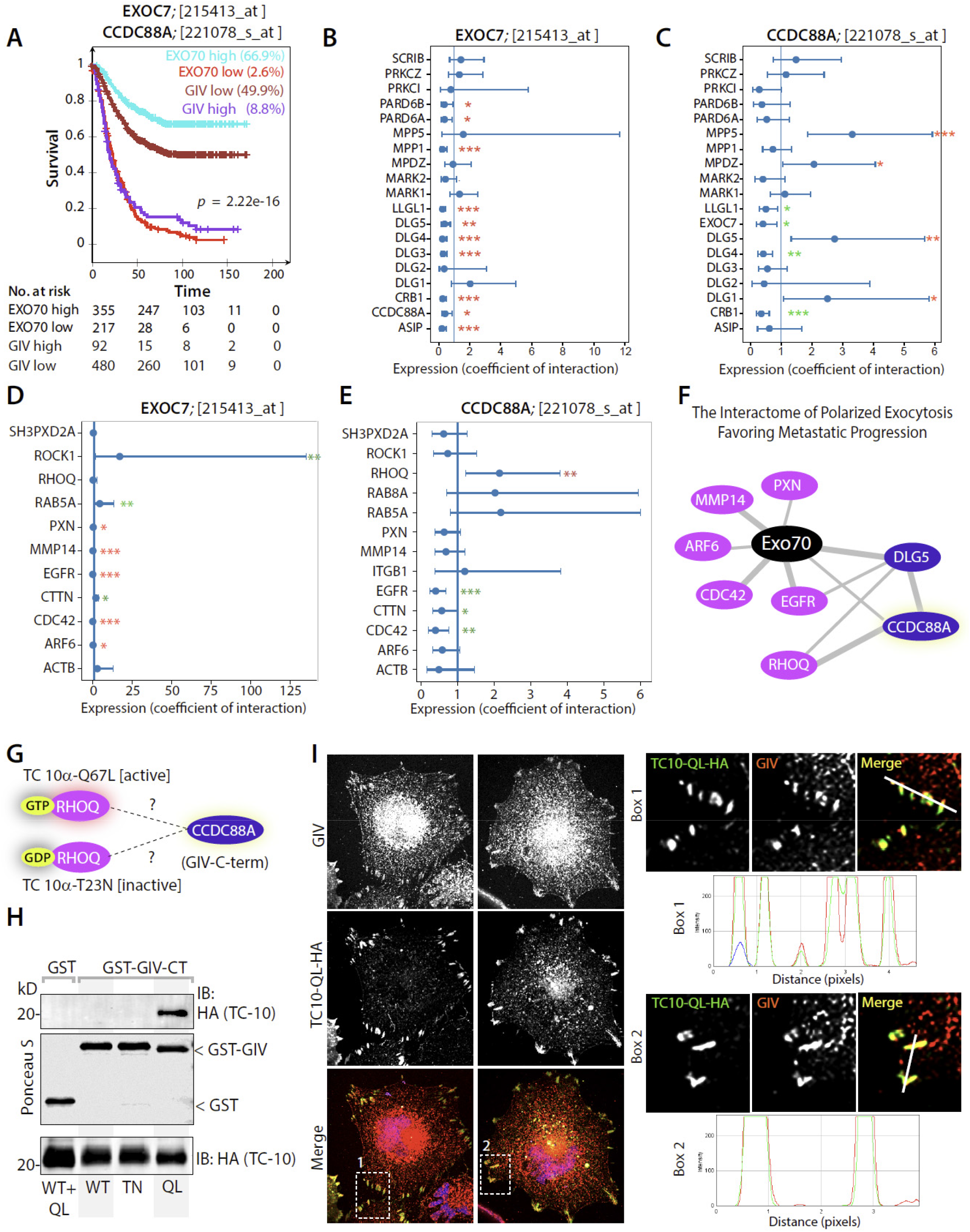
Polarity scaffold GIV (CCDC88A), matrix metalloprotease MT1-MMP and exocyst complexes synergize during metastasis. **A)** An overlay of Kaplan–Meier curves for metastasis-free survival over time among 572 patients [3 independent cohorts, pooled (Bos et al., 2009; Minn et al., 2005; Wang et al., 2005)] who did not receive adjuvant therapy segregated into high vs. low levels of exo70 (EXOC7) or GIV (CCDC88A) in primary tumors. **B-C)** Statistical interaction (synergy between variables) is measured in the Cox proportional hazards regression model for EXOC7 (exo70; **B**) or CCDC88A (GIV; **C**) and all mammalian polarity scaffold proteins, including CCDC88A (GIV). Coefficient of the interaction term in Cox regression models is plotted with 95% confidence interval that demonstrates the significance of the statistical interaction. *p* values: ‘**’ 0.01 ‘*’ 0.05. Red and green asterisks denote shorter or longer metastasis-free survival, respectively. **D-E)** Statistical interaction (synergy between variables) is measured in the Cox proportional hazards regression model as in B-C for EXOC7 (exo70; **D**) or CCDC88A (GIV; **E**) and mammalian proteins known to be involved in or influence the exocytosis of MT1-MMP. Red and green asterisks denote shorter or longer metastasis-free survival, respectively. **F)** Schematic summarizing only those interactions with EXOC7 that reduced metastasis-free survival. Purple = genes that encode polarity proteins; pink = others. Network-based prediction of RHOQ (TC10) as the RHO GTPase counterpart in mammals that synergistically interacts with CCDC88A (GIV). **G-H)** *<u>Top</u>*: Schematic summarizing the constructs used in **H** to assess nucleotide state-dependent interaction between TC10 and GIV. GST pulldown assays were carried out using lysates of COS7 cells as source of TC10-HA WT, or constitutively active (QL) or inactive (TN) mutants with GST or GST-GIV-CT. Bound proteins were assessed with anti-HA mAb. **I)** MDA MB231 cells exogenously expressing TC10-WT-QL (green) were fixed and stained for endogenous GIV (red). Representative images of transfected cells are shown. Red and green panels are shown in grey scale. Insets (Box1, 2) were analyzed for colocalization by line scans to generate RGB plots using ImageJ.

Based on all gene pairs tested, we computed the interactome for Exo70 that shortens metastasis-free survival (**Fig 6F**). The interactome revealed two major takeaways: 1) Polarized exocytosis during metastasis may require Exo70 to synergize with both GIV and DLG5. Because GIV binds DLG5 and their interaction regulates the formation of invadopodia (Ke et al., 2017), and DLG5 in turn engages with tSNARE syntaxin-4 (Nechiporuk et al., 2007) to enable the exocytosis (Wang et al., 2014), it is possible that DLG5 also impacts polarized exocytosis. 2) The interactome predicted that TC10, a close relative of Cdc42 is a likely candidate Rho-superfamily GTPase that may impact GIV’s ability to serve as the polarity scaffold during the exocytosis of MT1-MMP.

We asked if GIV’s C-terminus interacts with TC10 and if such interaction is regulated by its nucleotide-bound (activity) state. Using previously validated TC10 constructs (Chiang et al., 2002) that mimic active (QL) or inactive (TN) conformations (**Fig 6G**) in pulldown assays with GST-GIV-CT we first confirmed that GIV exclusively binds active, but not the inactive GTPase (**Fig 6H**). To translate the relevance of these *in vitro* findings to cells, we tested the impact of TC10 on the localization of endogenous full length GIV, as determined by confocal immunofluorescence on fixed cells. Transfection of active QL mutant, but not WT TC10 was sufficient to enhance the localization of GIV to focal adhesion and/or invadosome-like peripherally located structures (**Fig 6I, S7**), where they colocalized (**Fig 6I**, RGB plots). These findings are in keeping with the fact that mammalian Exo70 has been shown to directly interact with active TC10 (Inoue et al., 2003) and that activation of TC10 has been linked with the formation and functions of the invadosome. Together, our findings suggest that TC10 may be the evolutionary counterpart that fulfils the role that cdc42p has been shown to perform alongside the Exo70p•Bem1p complex in yeast (Liu and Novick, 2014).

## CONCLUSIONS

The major discovery we report herein is the mechanism and consequences of a direct interaction between GIV and Exo70, and its impact on polarized exocytosis of matrix metalloprotease MT1-MMP. Using selective single point mutants of GIV that are incapable of binding Exo70, we chart the two major consequences of this GIV•Exo70 interaction--First, the GIV•Exo70 interaction impacts our understanding of exocytosis in mammalian cells. Findings reveal that GIV imparts polarization to the process of exocytosis. How does GIV do so? Previously, we showed that GIV’s ability to activate trimeric GTPase Gαi is critical for its ability to enhance insulin-stimulated exocytosis of glucose transporter, GLUT4-storage vesicles (GSVs)(Lopez-Sanchez et al.,2015b), which in part accounts for GIV’s ability to maintain insulin sensitivity in skeletal muscle cells (Ma, 2015). Others have demonstrated that the same G protein regulatory module is essential for GIV’s ability to activate Par3/6/aPKC polarity complexes (Sasaki et al., 2015). Because the Par3/6/aPKC complexes are highly versatile in their ability to orchestrate all forms of cell polarity, e.g., front-rear, apical-basal, planar cell, and polarity during migration (Suzuki and Ohno, 2006), and positive feedback loop signals are essential for the amplification of stochastically arising clusters of polarity factors that create and maintain such forms of polarity (McCaffrey and Macara, 2012), we propose that GIV enables polarized exocytosis in a bipartite manner: It’s ability to bind Exo70 imparts polarity, and its ability to activate the Par3/6/aPKC polarity complexes *via* G protein intermediates (i.e., ‘free’ Gβ□ dimers) serves as the positive feedback loop. Furthermore, GIV has been shown to use‘free’ Gβ□ intermediates to activate Rho-GTPases (Leyme et al., 2015) and to terminate Arf-GTPase signaling on endomembranes (Lo et al., 2015), which may serve as additional positive and negative feedback loops. Such feedback loops have been shown to be critical for imparting robustness to the process of polarized exocytosis in yeast (Howell et al., 2012): A positive feedback occurs wherein Cdc42p•GTP at the membrane can recruit the cytoplasmic Bem1p complex, which generates more Cdc42p•GTP from local Cdc42p•GDP by GEF-catalyzed exchange; a negative feedback loop that enables dispersal of the polarity complexes by affecting the duration of activation of Cdc42p•GTP or phosphoinhibition of Bem1p. Whether GIV is at the cross-roads of these regulatory loops with the Rho-GTPase TC10 or Rab/Arf-GTPases, and what might be the exact mechanism(s) that impart regulatory control and robustness to the mammalian polarized exocytic machinery remains unknown. Regardless, GIV’s multi-modular makeup (see **Fig 1A**), its functional diversity (Aznar et al., 2016a), its growing catalog of phosphomodifications (including the “Y” within the Exo70-binding DFY^1743^D motif (see **Fig 3**); phosphosite.org), and its ability to integrate signals at the cross-roads of diverse types of signaling pathways [reviewed in (Ghosh et al., 2017)] tempts us to speculate that its association with Exo70 may impose intricate regulatory control over the mammalian polarized exocytic machinery making it more precise, context-dependent, versatile, and adapted to the needs of multi-cellular organisms. In keeping with this assumption, it is noteworthy that the yeast polarity scaffold Bem1p has no known homologues in any multicellular organisms (Liu and Novick, 2014), whereas GIV (Coleman et al., 2016) and its associated aPKC/Par3/6 complexes (Thompson, 2013) are absent in yeast and co-evolved only in worms and flies and highly conserved in mammals. Thus, our findings reveal the extent of evolutionary divergence in the polarized exocytic machinery from yeast to mammals (see legend, **Fig 7**); although the octameric exocyst complex remained constant, each component in yeast is replaced by its functional counterpart in mammals, beginning with an abrupt change in the polarity determinant coinciding with the onset of multicellular life forms. While our findings suggest that the GIV•Exo70 co-complex may constitute the core component that imparts polarity to exocytosis, these may not be sufficient. It is noteworthy that just like Bem1p is not the only component affecting Exo70 localization in yeast (PI_4,5_P_2_, Cdc42, Rho1 and actin all play significant roles), GIV may not be the only component affecting polarized exocytosis in mammalian cells’ and Par/aPKC complexes, TC10, actin and others may contribute significantly.

**Figure 7.**
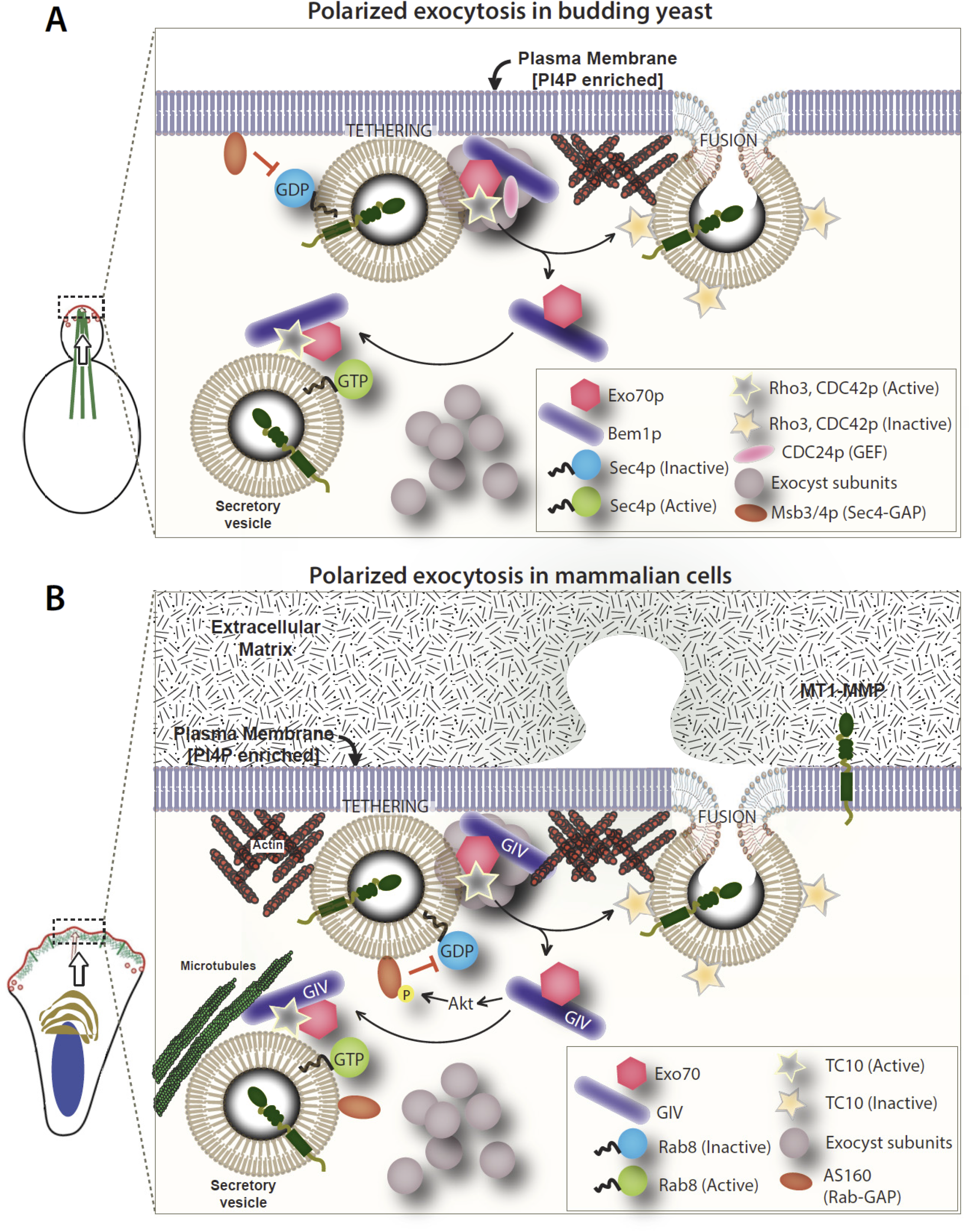
Components of polarized exocytosis display an evolutionary upgrade from yeast to mammals. Schematics summarizing the components of exocytosis in yeast (**A**) and mammals (**B**). Mammalian counterparts of all yeast components are represented using the same symbols. While the octameric exocyst complex is conserved, the polarity scaffolds Bem1p and GIV are divergent, alongside the previously annotated key components of the exocytic machinery. While sec4p in yeast is replaced by Rabs in mammals, the sec4p GAP (Msb2/3p) is replaced by the corresponding Rab-GAP, AS160, which has been previously shown to be activated by the GIV-dependent Class 1 PI3K»Akt signaling cascade (Ma et al., 2015).

Second, the GIV•Exo70 interaction impacts our understanding of the pro-metastatic role of GIV in diverse cancers and how it impacts key tumor cell behaviors that are dependent on proteolysis. Because GIV modulates integrin/FAK signaling *via* G protein intermediates (Leyme et al., 2016; Leyme et al., 2015; Lopez-Sanchez et al., 2015b), and because integrins and MT1-MMPs regulate each other’s activity, trafficking and signaling to integrate events in the extracellular milieu (Baciu et al., 2003; Das et al., 2017; Galvez et al., 2002; Gonzalo et al., 2010; Shi and Sottile, 2011; Zigrino et al., 2001), it is possible that GIV’s C-terminus may help in such process of integration. But to begin to entertain that possibility or test it, an in-depth insight into how GIV binds integrins, and how such binding impacts the GIV•Exo70 interaction is required first. Another issue that remains unexplored here is how the newly discovered GIV•Exo70 complex may be dysregulated in cancers. Increased tyrosine-based signaling and hyperphosphorylation of GIV at Y1743 may impact the GIV•Exo70 interaction directly. Alternatively, because both GIV (Ghosh et al., 2010; Puseenam et al., 2009) and Exo70 (Lu et al., 2013) have been shown to have alternative isoforms, pre-mRNA splicing may derail the GIV•Exo70 interaction or alter its functions indirectly. For example, in the case of Exo70, unlike the epithelial isoform which inhibits tumor metastasis, a mesenchymal isoform aids in tumor metastasis in an exocytosis-independent manner (Lu et al., 2013); the latter promotes tumor cell invasion by interacts with the Arp2/3 complex and stimulating actin branching in lamelipodia and invadopodia. Because GIV is a *bona-fide* actin remodeler that impacts actin dynamics (Enomoto et al., 2005; Jiang et al., 2008), it is possible that the GIV•(mesenchymal)Exo70 complex may collaboratively enhance these actin-based processes.

In conclusion, this work not just sheds light into GIV’s ability to enhance a key process that aids metastasis (i.e., polarized exocytosis of matrix metalloproteases), but also reveals how the modular makeup and functional diversity of GIV imparts the process of exocytosis features beyond polarity; it ‘upgrades’ the process by adding layers of regulatory control by diverse signaling pathways.

## ACKNOWLEDGEMENTS

This paper was supported by NIH CA238042, AI141630, CA100768 and CA160911 (to P.G). C.R was supported by an NCI/NIH-funded Cancer Therapeutics Training (CT^2^) Training Program (T32 CA121938) and a NIDDK/NIH-funded training grant (T32 DK007202). We thank Ying Dunkel and Nina Sun, for technical support in this work.

## AUTHOR CONTRIBUTIONS

C.R, N.R, I.L, and P.G designed, executed and analyzed most of the experiments in this work. P.N. provided critical tools (Bem1p and Exo70p) and guidance on experimental design. D.S. carried out multivariate analysis on the impact of gene expression on metastatic progression. C.R and P.G conceived the project, wrote materials and methods and edited the manuscript. P.G wrote the manuscript.

