## Supplementary Online Materials for "GIV/Girdin and Exo70 Constitute the Core of the Mammalian Polarized Exocytic Machinery"

### INVENTORY OF SUPPLEMENTARY MATERIALS


- **STAR METHODS**
- **SUPPLEMENTARY FIGURES AND LEGENDS**
- **SUPPLEMENTARY MOVIES AND LEGENDS**
- **REFERENCES CITED**

**STAR Methods**

- **KEY RESOURCE TABLE**
- **CONTACT FOR REAGENT AND RESOURCE SHARING**
- **EXPERIMENTAL MODEL AND SUBJECT DETAILS**
  - Cell Lines (Cos7, MDA-MB-231)
- **METHOD DETAILS**
  - Plasmid constructs
  - Protein Expression and Purification
  - Cell culture, transfection, lysis, and quantitative immunoblotting
  - Whole Cell Immunofluorescence and TIRF microscopy
  - Generation of stable cell lines
  - In Vitro Pulldown and Co-immunoprecipitation (Co-IP)
  - Gelatin Degradation Assay
  - Adhesion Assay
  - Transwell Invasion Assay
  - Analysis of gene expression data
- **QUANTIFICATION AND STATISTICAL ANALYSIS**
  - Statistical Analysis
  - Replications
- **DATA AND SOFTWARE AVAILABILITY**

### Key Resource Table:

| REAGENT or RESOURCE | SOURCE | IDENTIFIER |
|---|---|---|
| **Antibodies** | | |
| **Rabbit monoclonal anti-pY1765 GIV** | Roche Spring Biosciences | 06974937001 Clone SP158 |
| **Rabbit polyclonal anti-GIV CT** | Santa Cruz Biotechnology | N/A |
| **Rabbit polyclonal anti-Gαi3 (C-10)** | Santa Cruz Biotechnology | N/A |
| **Rabbit polyclonal anti-β-tubulin** | Santa Cruz Biotechnology | sc-9104 |
| **Mouse monoclonal anti-GAPDH** | Santa Cruz Biotechnology | sc-365062 |
| **Mouse monoclonal anti-HIS** | GenScript | A00186-100 |
| **Mouse monoclonal anti-GST** | GenScript | A00865 |
| **Mouse monoclonal anti-HA** | | |
| **Rabbit monoclonal anti-cortactin** | Santa Cruz Biotechnology | N/A |
| **Goat anti-Rabbit IgG (680)** | LI-COR Biosciences | 926-68071 |
| **Goat anti-Rabbit IgG, Alexa Fluor 594 conjugated** | ThermoFisher Scientific | A11072 |
| **Goat anti-Rabbit IgG, Alexa Fluor 647 conjugated** | ThermoFisher Scientific | A27040 |
| **Goat anti-Mouse IgG (800)** | LI-COR Biosciences | 926-32210 |
| **Goat anti-Mouse IgG, Alexa Fluor 488 conjugated** | Thermo Fisher Scientific | A11017 |
| **Biological Samples and Cell Lines** | | |
| **Cos7** | ATCC | CRL-1651 |
| **MDA-MB-231** | ATCC | HTB-26 |
| **Chemicals, Recombinant Proteins, and Plasmids** | | |
| **DAPI (4',6-Diamidino-2-Phenylindole, Dilactate)** | Thermo Fisher Scientific | D3571 |
| **Collagen I** | BD Biosciences | 354249 |
| **Poly-D-lysine** | Millipore Sigma | A-003-E |
| **Mirus TranIT LT1** | Mirus Bio LLC | MIR2300 |
| **G418** | Cellgro | A-1720 |
| **Puromycin** | Life Technologies | A1113803 |
| **Porcine Gelatin** | | |
| **Paraformaldehyde 16%** | Electron Microscopy Biosciences | 15710 |
| **Phalloidin 594** | Thermo Fisher Scientific | A12381 |
| **Prolong Gold** | Thermo Fisher Scientific | P10144 |
| **Prolong Glass** | Thermo Fisher Scientific | P36980 |
| **GST- Exo70 (human)** | *Xiong X. et al., Mol Biol Cell., 2012.*<br>*--Kun Ling* | N/A |
| **Exo-70-HA (human)** | *Sakurai-Yageta et al., 2008.*<br>*--Phillipe Chavrier* | N/A |
| **pGEX-4T-GIV-CT-WT (a.a. 1623-1870)** | *Garcia-Marcos et al., 2009* | N/A |
| **pET-28b-GIV-CT-WT (a.a. 1660-1870)** | *Garcia-Marcos et al., 2009* | N/A |
| **pET-28b-GIV-CT-PGxA (a.a. 1660-1870)** | This paper | N/A |
| **pET-28b-GIV-CT-AAxA (a.a. 1660-1870)** | This paper | |
| **pET-28b-GIV-CT-PAxF (a.a. 1660-1870)** | This paper | |
| **pET-28b-GIV-CT-AAxF (a.a. 1660-1870)** | This paper | |
| **pET-28b-GIV-CT-PGRF (a.a. 1660-1870)** | This paper | |
| **pET-28b-GIV-CT-DFYR (a.a. 1660-1870)** | This paper | |
| **pET-28b-GIV-CT-AFYD (a.a. 1660-1870)** | This paper | |
| **pET-28b-GIV-CT-DFYA (a.a. 1660-1870)** | This paper | |
| **pET-28b-GIV-CT-DFAD (a.a. 1660-1870)** | This paper | |
| **pET-28b-GIV-CT-AFYA (a.a. 1660-1870)** | This paper | |
| **pET-28b-GIV-CT-DFYDL (a.a. 1660-1870)** | *Lin et. al, MBoC 2014* | |
| **CMV14-p3X FLAG-GIV WT (full length)** | This paper | N/A |
| **CMV14-p3X FLAG-GIV P1742A (full length)** | This paper | N/A |
| **CMV14-p3X FLAG-GIV AAxA (full length)** | This paper | N/A |
| **pGEX4T2-Exo70p full length** | *Liu and Novick, JCB 2014*<br>*--Peter Novick* | N/A |
| **pGEX4T2-Exo70p Δ62N (Exo70Δ1-62)** | *Liu and Novick, JCB 2014* | N/A |
| **pGEX4T2-Exo70p Δ dA (Exo70 Δ 1-140)** | *Liu and Novick, JCB 2014*<br>*--Peter Novick* | N/A |
| **pGEX4T2-Exo70p Δ dB (Exo70 Δ 153-302)** | *Liu and Novick, JCB 2014*<br>*--Peter Novick* | N/A |
| **pGEX4T2-Exo70p Δ dAB (Exo70 Δ 1-338)** | *Liu and Novick, JCB 2014*<br>*--Peter Novick* | N/A |
| **pGEX4T2-Exo70p Δ dC (Exo70 Δ 347-514)** | *Liu and Novick, JCB 2014* | N/A |

| | | |
|---|---|---|
| | *--Peter Novick* | |
| **pGEX4T2-Exo70p Δ dD (Exo70 Δ 542-623)** | *Liu and Novick, JCB 2014 --Peter Novick* | N/A |
| **pGEX4T2-Exo70p, M26 (pGEX4T2Exo70S196F/L246S/K354A/K355A/E488A/K489A/E505A/R506A)** | *Liu and Novick, JCB 2014 --Peter Novick* | |
| **pGEX4T2-Exo70p, M30 (pGEX4T2-Exo70L246S/K354A/K355A/E488A/K489A/E505A/R506A)** | *Liu and Novick, JCB 2014 --Peter Novick* | |
| **pGEX4T2-Bem1** | *Liu and Novick, JCB 2014 --Peter Novick* | |
| **pET21a-Bem1** | *Liu and Novick, JCB 2014 --Peter Novick* | |
| **MT1-MMP–mCherry** | *Sakurai-Yageta et al, JCB 2008 --Phillipe Chavrier* | |
| **MT1-MMP-pHLuorin** | *Lizarraga et al Cancer Res. 2009 – Phillipe Chavrier* | |
| **Software** | | |
| **ImageJ** | National Institute of Health | https://imagej.net/Welcome |
| **Prism** | GraphPad | https://www.graphpad.com/scientific-software/prism/ |
| **LAS-X** | Leica | www.leica-microsystems.com/products/microscope-software/p/leica-las-x-ls |

### Methods

#### Plasmid constructs

Cloning of GIV-CT (aa 1660–1870) into pET28b (His-GIV CT) and GIV-CT (aa 1623-1870) were described previously (Garcia-Marcos et al., 2012). pGEX4T2Exo70p, pGEX4T2-Exo70Δ62N (Exo70Δ1-62), pGEX4T2-Exo70 Δ dA (Exo70 Δ 1-140), pGEX4T2-Exo70 Δ dB (Exo70 Δ 153-302), pGEX4T2-Exo70 Δ dAB (Exo70 Δ 1-338), pGEX4T2-Exo70 Δ dC (Exo70 Δ 347-514), pGEX4T2-Exo70 Δ dD (Exo70 Δ 542-623), pGEX4T2-Exo70M26, pGEX4T2-Exo70M30, pGEX4T2-Bem1, and pET21a-Bem1 plasmids were generously obtained from Peter Novick (UCSD, California). MT1-MMP–mCherry and MT1-MMP-pHLuorin plasmids were generously shared by Phillipe Chavrier. For mammalian expression, RNA interference–resistant (shRNA rest) GIV was cloned into p3XFLAG-CMVTM-14 plasmid (GIV-FLAG) as described previously (Garcia-Marcos et al., 2009). GIV-FLAG and His-GIV-CT mutants (detailed in table) were generated by site-directed mutagenesis using a QuikChange kit (Stratagene) and specific primers (sequence available upon request) as per the manufacturer's protocols (Garcia-Marcos et al., 2009; Lin et al., 2011). Primer sequences are available upon request. shRNA 3–untranslated region for GIV (GIV shRNA:CCGGGCTTTCATT-ACCAGCTCTGAACTCGAGTTCAGAGCTGGTAATGAAAGCTTTTTTG) was cloned into pLKO.1 (TRCN0000130452) or control vector TRC1.5-pLKO.1-puro. GST-Exo70 (Xiong et al., 2012) was obtained from Kun Ling (Mayo Clinic). Exo70-HA (Sakurai-Yageta et al., 2008) was obtained from Philippe Chavrier (Institut Curie, France).

#### Protein expression and purification

Both GST and His-tagged proteins were expressed in E. coli stain BL21 (DE3) and purified as previously described (Garcia-Marcos et al., 2009; Ghosh et al., 2008). Briefly, cultures were induced using 1mM IPTG overnight at 25°C. Cells were then pelleted and resuspended in either GST lysis buffer (25 mM Tris-HCl, pH 7.5, 20 mM NaCl, 1 mM EDTA, 20% (vol/vol) glycerol, 1% (vol/vol) Triton X-100, 2×protease inhibitor cocktail) or His lysis buffer (50 mM $NaH_2PO_4$ (pH 7.4), 300 mM NaCl, 10 mM imidazole, 1% (vol/vol) Triton X-100, 2X protease inhibitor cocktail). Cells were lysed by sonication, and lysates were cleared by centrifugation at 12,000 X g at 4°C for 30 mins. Supernatant was then affinity purified using glutathione-Sepharose 4B beads (GE Healthcare) or HisPur Cobalt Resin (Thermo Fisher Scientific), followed by elution, overnight dialysis in PBS, and then storage at -80°C.

His-GIV-CT fusion construct was expressed in E. coli strain BL21 (DE3) and purified as described previously. Briefly, bacterial cultures were induced overnight at 25°C with 1 mM isopropyl β-d-1-thiogalactopyranoside (IPTG). Pelleted bacteria from 1 l of culture were resuspended in 10 ml of His lysis buffer (50 mM NaH2PO4, pH 7.4, 300 mM NaCl, 10 mM imidazole, 1% [vol:vol] Triton X-100, 2X protease inhibitor cocktail [Complete EDTA-free, Roche Diagnostics, CA, USA]). After sonication (3 × 30 s), lysates were centrifuged at 12,000 × g at 4°C for 20 min. Solubilized proteins were affinity purified on HisPur Cobalt Resin (Pierce, IL, USA). Proteins were eluted, dialyzed overnight against PBS, and stored at −80°C.

***Cell culture, transfection, lysis, and quantitative immunoblotting***

Unless mentioned otherwise, cell lines used in this work were cultured according to American Type Culture Collection (ATCC) guidelines or guidelines previously published for each line Cos7, HeLa and MDA-MB-231 cells were obtained from American Type Culture Collection (ATCC). Transfection, lysis, and immunoblotting were carried out exactly as described before (Aznar et al., 2016; Lopez-Sanchez et al., 2015) Cells were transfected using Mirus LT1 following the manufacturers' protocols. For assays involving serum starvation, serum concentration was reduced to 0% FBS overnight.

Whole-cell lysates were prepared after washing cells with cold PBS before resuspending and boiling them in sample buffer. Lysates used as a source of proteins in pull-down assays were prepared by resuspending cells in lysis buffer (20 mM HEPES, pH 7.2, 5 mM Mg acetate, 125 mM K acetate, 0.4% Triton X-100, and 1 mM dithiothreitol supplemented with sodium orthovanadate [500 μM], phosphatase [Sigma-Aldrich, MO, USA], and protease [Roche, USA] inhibitor cocktails), after which they were passed through a 28-gauge needle at 4°C and cleared (10,000 × g for 10 min) before use in subsequent experiments.

For immunoblotting, protein samples were separated by SDS-PAGE and transferred to polyvinylidene fluoride membranes (Millipore Sigma, MO, USA). Membranes were blocked with PBS supplemented with 5% nonfat milk (or with 5% BSA when probing for phosphorylated proteins) before incubation with primary antibodies. In some instances, the samples were separated on a 12% SDS PAGE and transferred to a nitrocellulose membrane (Bio-Rad, CA, USA) using TransBlot-Turbo (Bio-Rad, CA, USA). The membrane was stained with Ponceau S to visualize baits, then washed and blocked with PBS with 0.1% Tween (PBS-T) and 0.5% BSA overnight at 4°C. Infrared imaging with two-color detection and quantification were performed using a Li-Cor Odyssey imaging system. Dilution of primary antibodies used were as follows: anti–GIV-CT, 1:500; anti-Gαi3, 1:333; anti-β tubulin, 1:1000; and anti–His, 1:500. All blots were visualized using LI-COR Odyssey imager, and band analysis was performed with Image Studio™ Lite 5.2 (LI-COR Biosciences, NE, USA). Figures were assembled for presentation using Photoshop (Adobe, San Jose, CA, USA) and Illustrator (Adobe, San Jose, CA, USA) software.

#### *In Vitro Pulldown and Co-immunoprecipitation (Co-IP)*

Purified GST-tagged proteins from E. coli were immobilized onto glutathione-Sepharose beads and incubated with binding buffer (50 mM Tris-HCl pH 7.4, 100 mM NaCl, 0.4% (v:v) Nonidet P-40, 10 mM MgCl2, 5 mM EDTA, 2 mM DTT, 1X Complete protease inhibitor) for 60min at room temperature. For GST-pulldown assays with recombinant proteins, the proteins were diluted in binding buffer and incubated with immobilized GST-proteins for 90min at room temperature. For binding with cell lysates, cells were lysed in cell lysis buffer (20 mM HEPES pH 7.2, 5 mM Mg-acetate, 125 mM K-acetate, 0.4% Triton X-100, 1 mM DTT, 500 µM sodium orthovanadate, phosphatase inhibitor cocktail (Sigma-Aldrich, MO, USA) and protease inhibitor cocktail (Roche Life Science)) using a 28G syringe, followed by centrifugation at 10,000Xg for 10min. Cleared supernatant was then used in binding reaction with immobilized GST-proteins for 4 hours at 4°C. After binding, bound complexes were washed four times with 1 ml phosphate wash buffer (4.3 mM Na2HPO4, 1.4 mM KH2PO4, pH 7.4, 137 mM NaCl, 2.7 mM KCl, 0.1% (v:v) Tween 20, 10 mM MgCl2, 5 mM EDTA, 2 mM DTT, 0.5 mM sodium orthovanadate). Bound proteins were then eluted through boiling at 100°C in Laemmli buffer (BIORAD, CA, USA).

#### *Whole-cell immunofluorescence*

Cells were fixed at room temperature with 3% PFA in PBS for 15 min, treated with 0.1 M glycine for 10 min, and subsequently permeabilized for 1 h (0.2% Triton X-100 in PBS) and blocked in PBS containing 1% bovine serum albumin (BSA) and 0.1% Triton X-100 as described previously (Lopez-Sanchez et al., 2014 blue right-pointing triangle). Primary and secondary antibodies where incubated for 1 h at room temperature in blocking buffer. ProLong Gold or Prolong Glass (Life Technologies) was used as mounting medium. Dilutions of antibodies used were as follows: HA, 1:500; phalloidin, 1:1000; cortactin, 1:500; DAPI, 1:2000; and secondary goat anti-rabbit (488), goat anti-mouse (594), and goat-anti-mouse or rabbit (647) Alexa-conjugated antibodies, 1:500. Images were acquired at room temperature with a Leica TCS SP8 with DMI8 microscope equipped with a PMT and HyD detectors and the LAS X (Leica) software using a 100× oil-immersion objective using a white light laser tunable to 488-, 561-, 635-, and 405-nm for excitation. The settings were optimized, and the final images scanned with line averaging of three scans. Images were further processed using Lightning deconvolution in the Leica LAS-X software package. All images were processed using ImageJ software and assembled for presentation using Photoshop and Illustrator software (Adobe, San Jose, CA). Images shown are representative of 90–95% cells that were evaluated across three independent experiments.

Cells were plated on gelatin-coated coverslips and allowed to adhere for 30 min. They were then fixed with 4% PFA and stained with antibodies against cortactin (1:100), HA (1:100) or direct fluorescence from MT1-MMP mCherry construct followed by secondary antibody incubation (AlexaFluor 488 and AlexaFluor 647 1:500). Coverslips were then mounted on Prolong Glass and allowed to cure for 5 days before imaging. Once images were acquired, they were deconvolved using LAS-X software and then processed using Image-J. All images were further processed in Image-J using the 3D surface plot plugin and line scans were done to generate RGB plots.

#### *Generation of stable cell lines*

ShRNA control and shRNA GIV MDA-MB-231, stable cell lines using Mission RNAi technology (Sigma-Aldrich, MO, USA) were generated by lentiviral transduction followed by selection with puromycin (2.5 μg/ml) a described previously (Midde et al., 2018). Depletion of endogenous GIV was confirmed by immunoblotting with GIV-CT rabbit antibody. Lentiviral packaging was performed in HEK293T cells by co-transfecting the shRNA constructs with psPAX2 and pMD2G plasmids (4:3:1 ratio, respectively), using Mirus LT1. The medium was changed after 24 h, and virus-containing medium was collected after 36–48 h and centrifuged and filtered through a 0.45-μm filter. psPAX2 and pMD2G plasmids were a generous gift from Christopher K. Glass (University of California, San Diego, La Jolla, CA). shRNA GIV MDA-MB-231 stable cell lines expressing p3xFLAG-CMV-14-GIV (GIV-3xFLAG) WT, GIV-PGxA and AAxA constructs were selected as previously described (Aznar et al., 2016) with the neomycin analogue G418 at 800 μg/ml. Expression of various GIV constructs were confirmed to be similar to levels of endogenous GIV in shRNA control cells by immuno-blotting with GIV-CT antibodies.

#### *Fluorescent gelatin degradation assay*

MDA-MB-231 cells were incubated for 5 h on FITC-conjugated cross-linked gelatin, and then fixed and stained for F-actin. Cells were imaged with the 63× objective of a Leica TCS SPE confocal microscope and controlled by LAS software. For quantification of degradation, the total area of degraded matrix in one field (black pixels) measured with the Threshold command of ImageJ was divided by the number of phalloidin-labeled cells in the field to define a degradation index, which was normalized to the degradation index of control cells set to 100. All experiments were performed in triplicate, and each experiment was repeated at least three times.

#### *Total internal reflection of fluorescence (TIRF)- Microscopy*

MDA-MB-231 cells transfected with pHLuorin-tagged MT1-MMP were plated on glass-bottom dishes coated with cross-linked unlabeled gelatin as previously described. TIRFM sequences were acquired on an inverted microscope (TE2000) equipped with a 100× TIRF objective (1.47 NA), a TIRF arm, an image splitter (DV; Roper Scientific) installed in front of the CCD camera and a temperature controller. pHLuorin was excited with 491-laser, (100 mW; Roper Scientific), both controlled for power by an acousto-optic tunable filter. Fluorescent emissions were selected with bandpass and longpass filters (Chroma Technology Corp.) and captured by a QuantEM EMCCD camera (Roper Scientific). To measure the subplasma membrane density of MT1-MMP endosomes, cells stably expressing MT1-MMP-pHLuorin were plated on a layer of unlabeled gelatin and imaged by TIRFM for an exposure time of 100 ms and with 1 pixel corresponding to 160 nm. The density of MT1-MMP-pHLuorin structures was evaluated by ImageJ and quantified using the particle analyzer application on ImageJ (National Institutes of Health, Bethesda, MD) exactly as outlined previously (Horzum et al., 2014). The threshold was set at 1.3× the cytoplasmic background.

***Transwell invasion assay***

Cell invasion was assessed using Biocoat Matrigel (Corning, NY, USA) inserts with 8-μm pores in 24-well plates. Cells were detached using trypsin/EDTA and resuspended in DMEM supplemented with 0.4% FBS. A total of 5 × 105 cells was loaded in the upper well in a volume of 300 μl, and the lower well was filled with 750 μl of DMEM with 10% FBS. The plates were incubated at 37°C for 24h before removing the remaining cell suspension. The invasion insert was placed in a clean well containing 4% PFA for 1 h at room temperature, stained with crystal violet for 1 h, and washed three times in PBS. Cells on the upper side of the filters were removed with cotton-tipped swabs, and the number of migrated cells on the bottom side of the filter was counted in five randomly chosen fields at 200× magnification and averaged. All experiments were performed in triplicate, and each experiment was repeated at least three times.

***Adhesion Assay***

12-well plates were coated with collagen, rinsed with PBS, and then blocked with 0.5% BSA for 1h at 37°C. Cells were harvested with trypsin/EDTA, seeded in the wells at 2 × 104 cells/well in 1,000 μl of DMEM with 0.4% FBS, and allowed to adhere for 30min at 37°C. Nonadherent cells were removed by gentle washing twice with PBS, and attached cells were fixed in 4% PFA for 15 min and then stained with 2.3% crystal violet (Sigma-Aldrich, MO, USA) for 10min. Cells were extensively washed, air dried, and then the cells imaged and counted using ImageJ. All experiments were performed in triplicate, and each experiment was repeated at least three times.

***Analysis of gene expression data***

Gene expression data from three different cohorts of patients with breast cancers (Bos et al., 2009; Minn et al., 2005; Wang et al., 2005) were collected from the National Center for Biotechnology Information (NCBI) Gene Expression Omnibus (GEO) website (Barrett, T. et al. 2005; Edgar et al., 2002). The dataset was prepared by pooling data from GSE2034, GSE2603 and GSE12276 and normalizing them together using Robust Multi-chip Average (RMA) algorithm. Patient survival data were carefully annotated for Kaplan-Meier analysis. To derive optimal cut-off values of gene expression levels, they are ordered from low to high and a rising step function was computed to define a threshold by StepMiner algorithm (Sahoo et al., 2007). Gene expression values were converted to high and low levels based on the StepMiner threshold. A noise margin of +/- 0.5 was used around the StepMiner threshold to provide relaxed estimates of the high/low values. A noise margin of +/- 0.5 around StepMiner threshold was used to soften or harden the actual threshold. Time-dependent survival probabilities are estimated with the Kaplan-Meier method (Kaplan EL, 1958) and compared using the log-rank test (Peto et al., 1977). Cox proportional-hazards regression models and life-tables (Cox, 1972) were used to test the statistical interaction between two different genes based on their association with survival outcome. High and low expression patterns for focal genes such as GIV (CCDC88A), Exo70 (EXOC7) and Dlg5 (DLG5) were compared individually using Kaplan-Meier analyses in R statistical software (R version 3.4.4 2018-03-15). Statistical interaction (synergistic effects) between polarity determinant genes (Lin et al., 2015) and Exo70 (EXOC7) were measured using interaction terms in Cox proportional hazards regression model on the above pooled breast cancer dataset with survival data. For example, coefficient of interaction terms was computed from Cox regression model for CCDC88A and EXOC7 as follows:

$$h(t) = h_0(t)\exp(a1 * CCDC88A + a2 * EXOC7 + a3 * CCDC88A * EXOC7)$$

where $h(t)$ is the hazard rate at time $t$ for an individual; $a1, a2,$ and $a3$ are the regression coefficients; $h_0(t)$ is the baseline hazard; two indicator variables $CCDC88A$ and $EXOC7$ (high expression = 1 else 0) that had only additive or interactive effects; $a3$ is the coefficient of interaction terms.

### Data analysis and other methods

All experiments were repeated at least three times, and results were presented either as one representative experiment or as average ± SD. Statistical significance was assessed with the Student's t test. Statistical significance between datasets with three or more experimental groups was determined using one-way analysis of variance (ANOVA) including a Tukey's test for multiple comparisons. For all tests, a p-value of 0.05 was used as the cutoff to determine significance [*p < 0.05, **p < 0.01, ***p < 0.001, ****p < 0.0001]. All experiments were repeated a least three times, and p-values are indicated in each figure. All statistical analysis was performed using GraphPad prism 8.

### Data and resource sharing

All data and constructs are available for sharing.

### SUPPLEMENTARY FIGURES AND LEGENDS

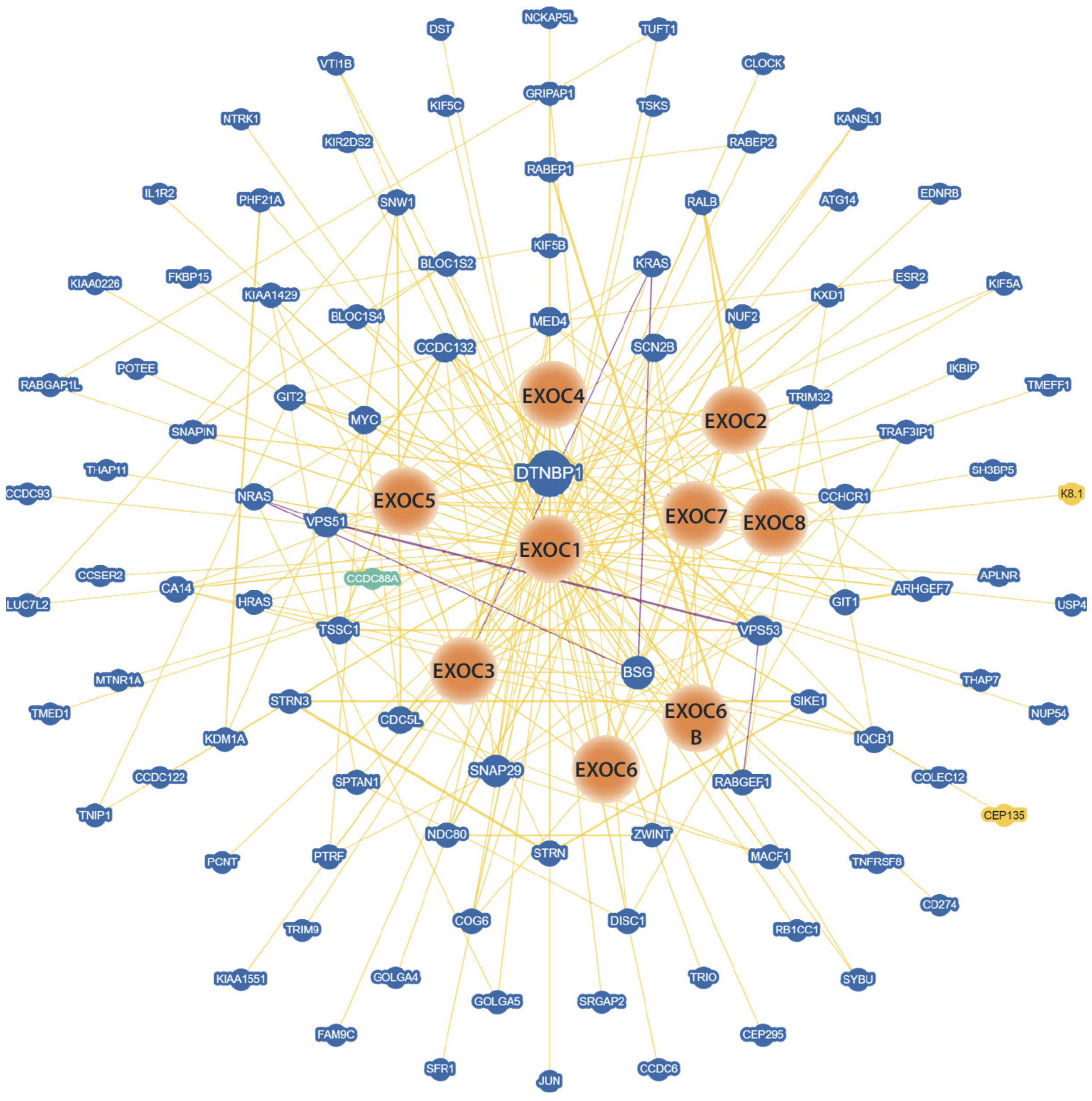


**Supplementary Figure 1 [related to Figure 1]**

**The exocyst interactome includes the polarity scaffold GIV/Girdin**. Visualization of the physical associations the subunits of the exocyst complexes using BioGRID (Version 3.5.178). CCDC88A (GIV/Girdin) is highlighted in green. Exocyst complex subunits are highlighted in orange. Blue nodes = associated protein from same species; yellow nodes = associated protein from another organism.

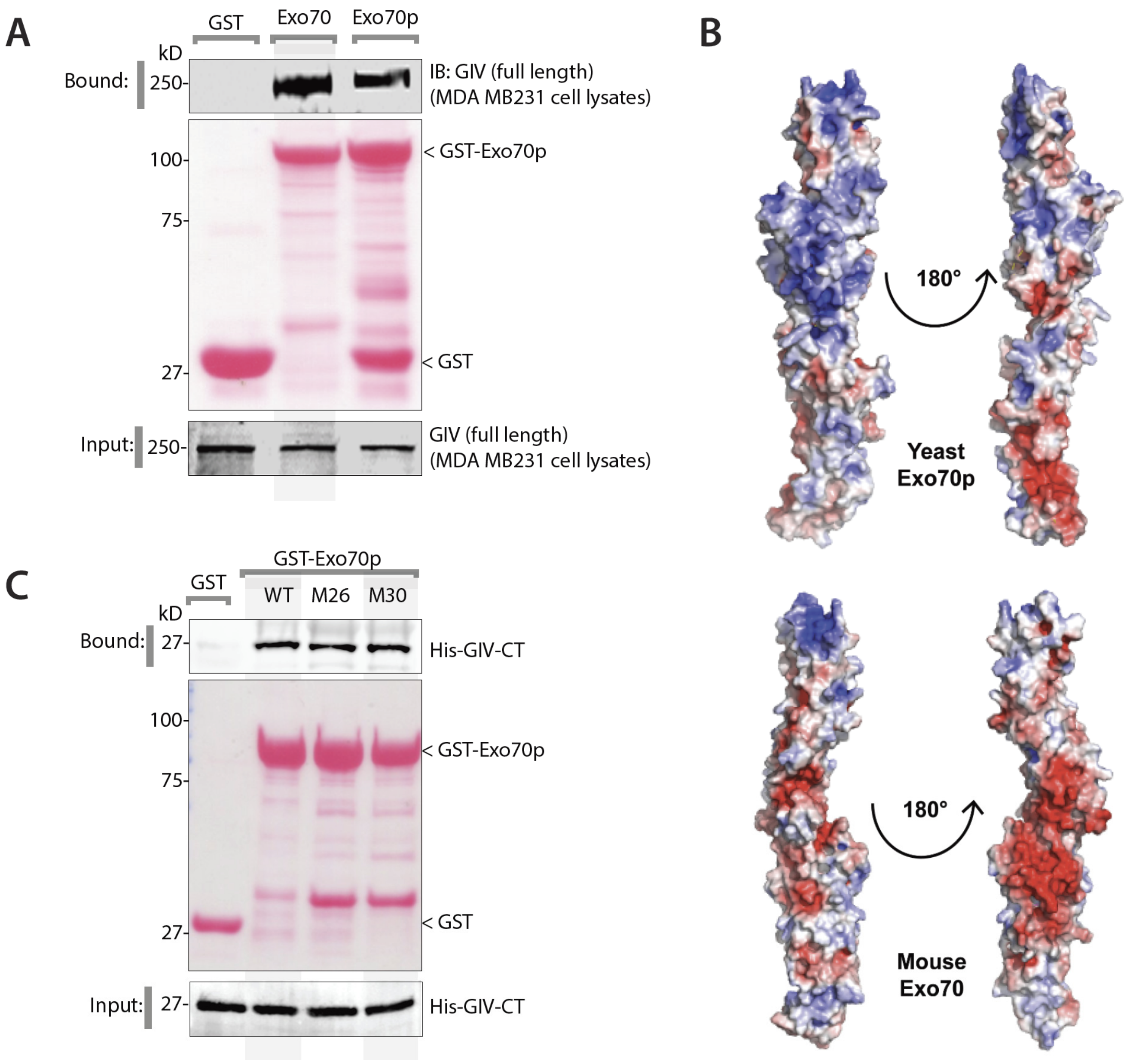


**Supplementary Figure 2 [related to Figure 1]**

**GIV binds Exo70 similar to Bem1p but contact residues on Exo70 differ.**

**(A)** Equal aliquots of lysates of MDA-MB-231 cells were used as source for full-length endogenous GIV in GST pulldown assays with GST, GST-Exo70p and GST-Exo70 proteins. Bound GIV was visualized by immunoblotting using anti-GIV-CT rabbit polyAb. Equal loading of GST proteins was confirmed by Ponceau S staining.

**(B)** Comparison of yeast Exo70p [PDB: 2B7M; (Hamburger et al., 2006)] and mouse Exo70 [PDB: 2PFT; (Moore et al., 2007)] structures, visualized using Molsoft ICM pro.

**(C)** Equal aliquots of recombinant His-GIV-CT (~ 3 µg) were used in GST pulldown assays with GST, GST-Exo70p-WT and two previously defined Bem1p-binding deficient complex mutants of GST-Exo70 (mutant M26 and M30) (Liu and Novick, 2014). Bound GIV was visualized by immunoblotting using anti-His mAb. Equal loading of GST proteins was confirmed by Ponceau S staining.

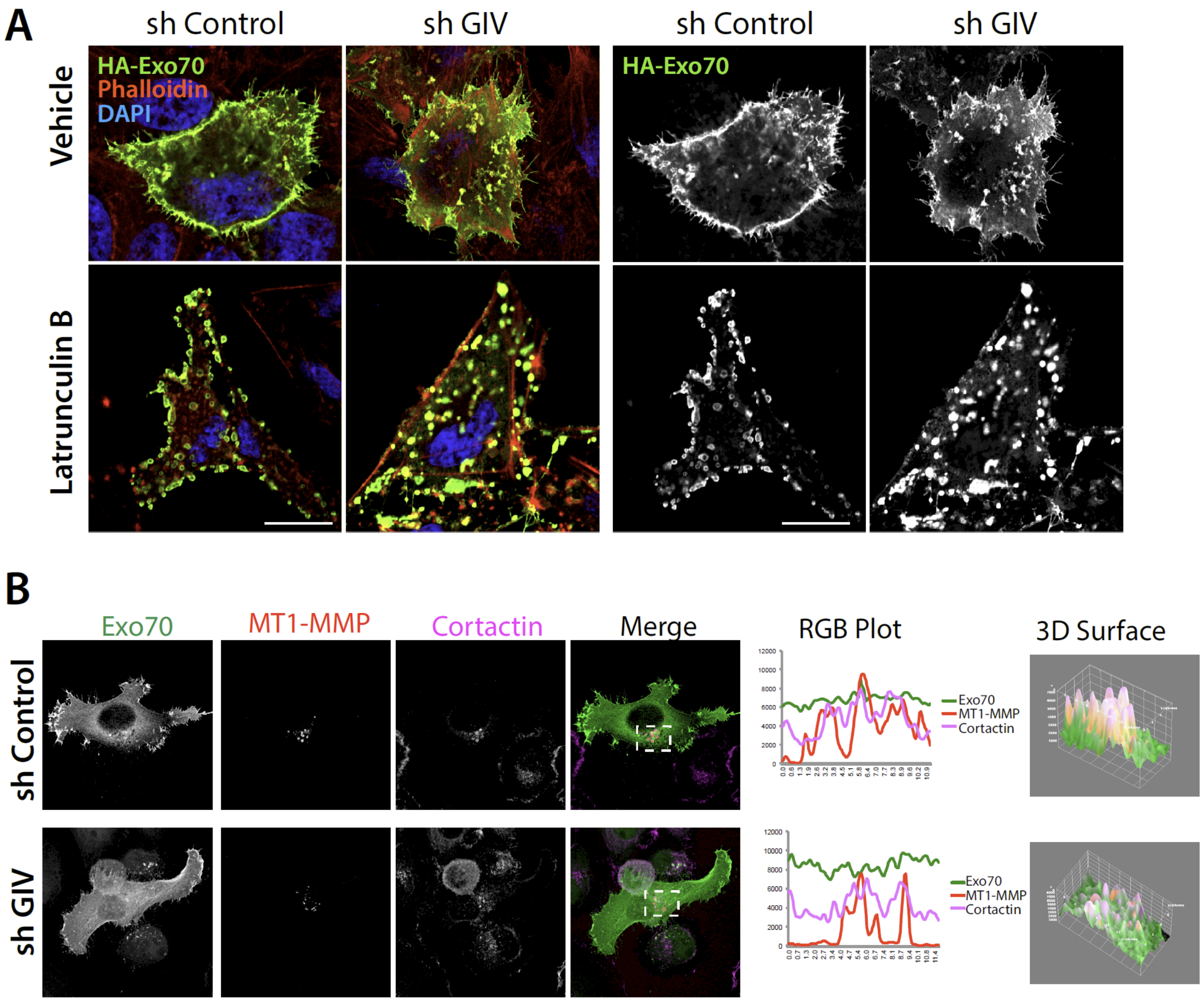


**Supplementary Figure 3 [related to Figure 2]**

**GIV is required for the localization of Exo70 and for the delivery of trans-membrane MT1-MMP cargo to the podosomes.**

**(A)** Control (sh Control) or GIV-depleted (sh GIV) MDA-MB-231 cells exogenously expressing HA-Exo70 were treated (lower panel) or not (vehicle) with a drug that depolymerizes actin cytoskeleton, Latrunculin B (25 µM, 24 h) prior to fixation with PFA. Fixed cells were stained for HA (green, Exo70), Phalloidin (red; actin) and DAPI (blue, nuclei) and analyzed by confocal microscopy. Representative deconvolved images are shown.

**(B)** Control (sh Control) or GIV-depleted (sh GIV) MDA-MB-231 cells exogenously co-expressing HA-Exo70 and MT1-MMP-mCherry plated on gelatin were fixed and stained for cortactin (a marker of cellular podosome). Representative deconvolved images are shown. Insets were analyzed by rendering 3D surface plots and line scans were taken to generate RGB plots using ImageJ.

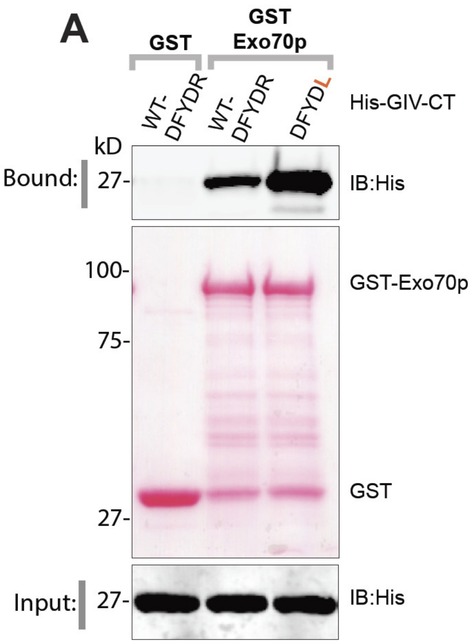


**B**

| GIV Mutants Tested | Binding to Exo70 |
|---|---|
| PGDAYDR$^{1745}$ | ---- |
| PADFYDR$^{1745}$ | - |
| AADFYDR$^{1745}$ | - |
| AADAYDR$^{1745}$ | ---- |
| PGRFYDR$^{1745}$ | ---- |
| PGDFYRR$^{1745}$ | + |
| PGAFYDR$^{1745}$ | - |
| PGDFYAR$^{1745}$ | ---- |
| PGDFADR$^{1745}$ | -- |
| PGAFYAR$^{1745}$ | -- |
| PGDFYDL$^{1745}$ | +++ |

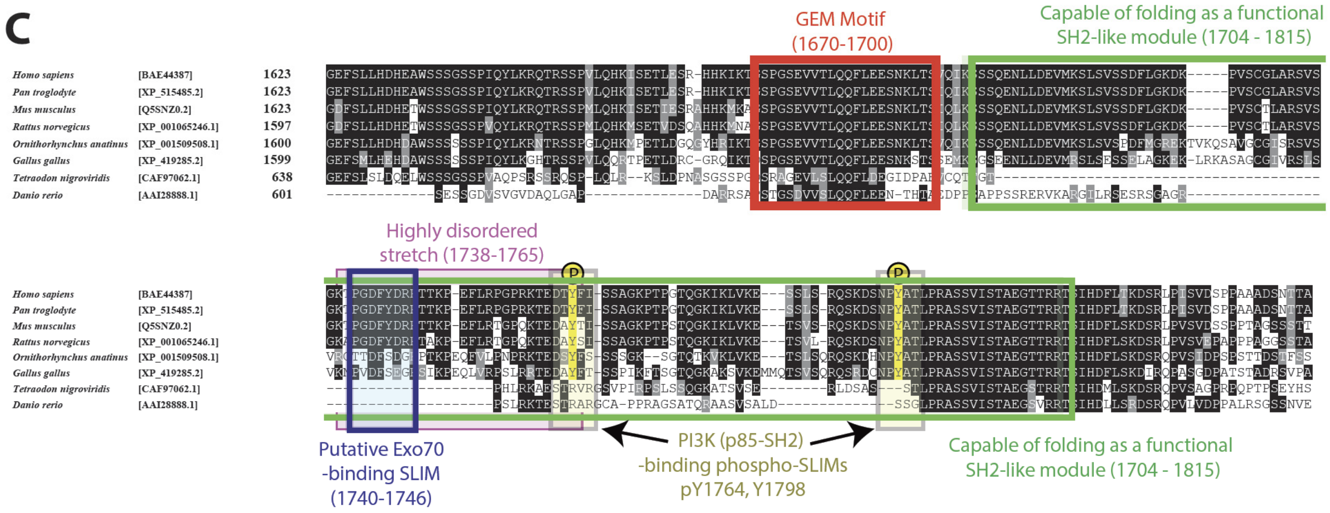


**Supplementary Figure 4 (related to Fig 3)**

**A short linear motif within the disordered C-terminus of GIV mediates binding to Exo70.**

**(A)** Equal aliquots of WT ($^{1741}$DFYDR$^{1745}$) or mutant ($^{1741}$DFYD**R»L**$^{1745}$) recombinant His-GIV-CT (~ 3 µg) were used in GST pulldown assays with GST and GST-Exo70p. Bound GIV was visualized by immunoblotting using anti-His mAb. Binding of the $^{1741}$DFYD**R»L**$^{1745}$ mutant GIV was consistently seen to be increased ~2-3 fold. Equal loading of GST proteins was confirmed by Ponceau S staining.

**(B)** Table summarizing the impact of various GIV mutations within and flanking the short linear motif $^{1739}$PGDFYDR$^{1745}$ on Exo70 binding, as assessed during *in vitro* GST pulldown assays (see **Fig 3**).

**(C)** Sequence alignment of GIV's C-terminus displaying a complete catalog of previously validated short linear motifs (SLIMS) and modules; the position of the newly identified Exo70-binding SLIM (aa 1740-1746) is shown in black. Because this SLIM is located within a highly disordered stretch (pink; aa 1738-1765; identified using InterPro [https://www.ebi.ac.uk/interpro/] and confirmed using MobiDB [http://mobidb.bio.unipd.it/]), it is likely to be impacted by conformational plasticity. Because multiple phosphosites flank this SLIM, it is likely that binding of GIV to Exo70 is regulated by post-translational modifications.

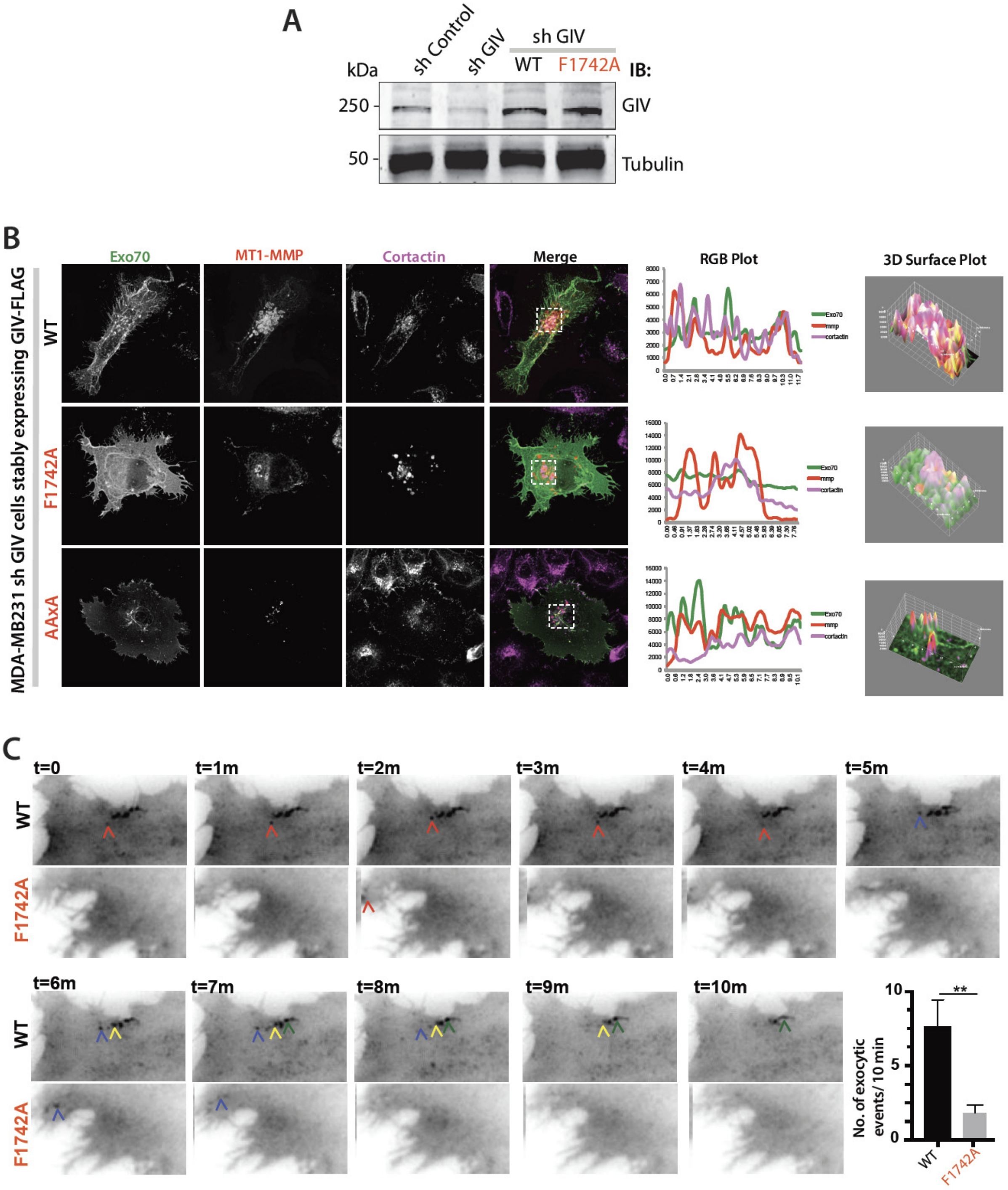


**Supplementary Figure 5 (related to Fig 4)**

**The Exo70•GIV interaction is required for the exocytic delivery of MT1-MMP to cellular podosomes.**

**(A)** Immunoblots of lysates of control (sh Control) and GIV-depleted (sh GIV) MDA MB231 cells stably expressing GIV-FLAG WT and Exo70-binding deficient F1742A mutant constructs.

**(B)** GIV-depleted (sh GIV) MDA-MB-231 cells stably expressing WT or mutant full length GIV proteins were transfected with HA-Exo70 (green) and MT1-MMP-mCherry (red) were fixed and stained for cortactin (a marker of cellular podosome). Representative deconvolved images are shown. Insets were analyzed by rendering 3D surface plots and line scans were taken to generate RGB plots using ImageJ.

**(C)** GIV-depleted (sh GIV) MDA-MB-231 cells stably expressing WT or mutant full length GIV proteins were transfected with MT1-MMP-pHLuorin were plated on a layer of unlabeled gelatin and imaged by TIRF microscopy. _Left:_ Still frames from a 10 min long movie (Supplementary movies 3, 5). _Bottom right:_ Bar graph shows the number of exocytic events that were encountered during the 10-min long span. Each colored arrowhead tracks one vesicle. N = ~ 5-10 cells/assay. Error bars = S.E.M. *p* values, **0.01.

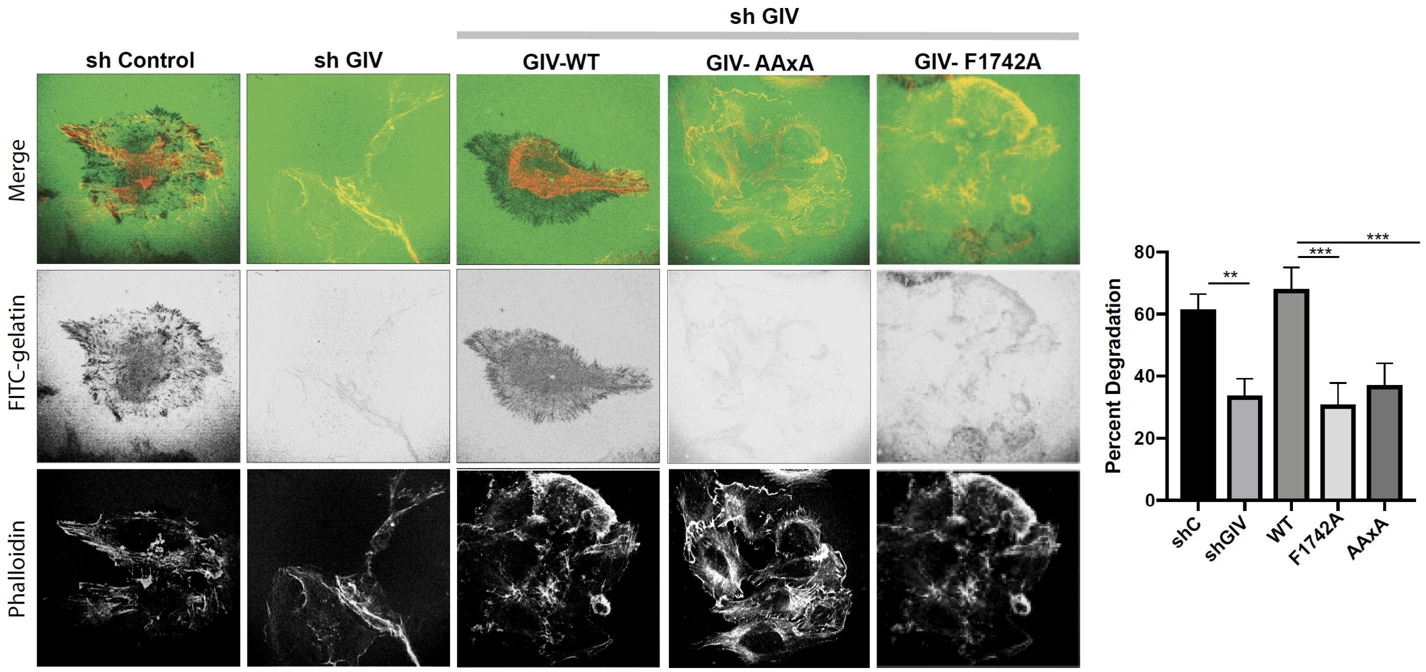


**Supplementary Figure 6 (related to Fig 4)**

**The Exo70•GIV interaction is required for the degradation of extracellular matrix at focal adhesions.**

*Left*: Control (sh Control), GIV-depleted (sh GIV) MDA-MB-231 cells and GIV-depleted cells stably expressing full length WT or Exo70-binding deficient mutant GIV proteins were plated for 5 h on FITC-conjugated cross-linked gelatin (green), and then fixed and stained for F-actin (Phalloidin; red). Representative images are shown. *Right*: Bar graphs display the % of cells that showed degradation of gelatin. N = ~100-200 cells /experiment x 3. Error bars = S.E.M. *p* values, * 0.05; **0.01; ***0.001.

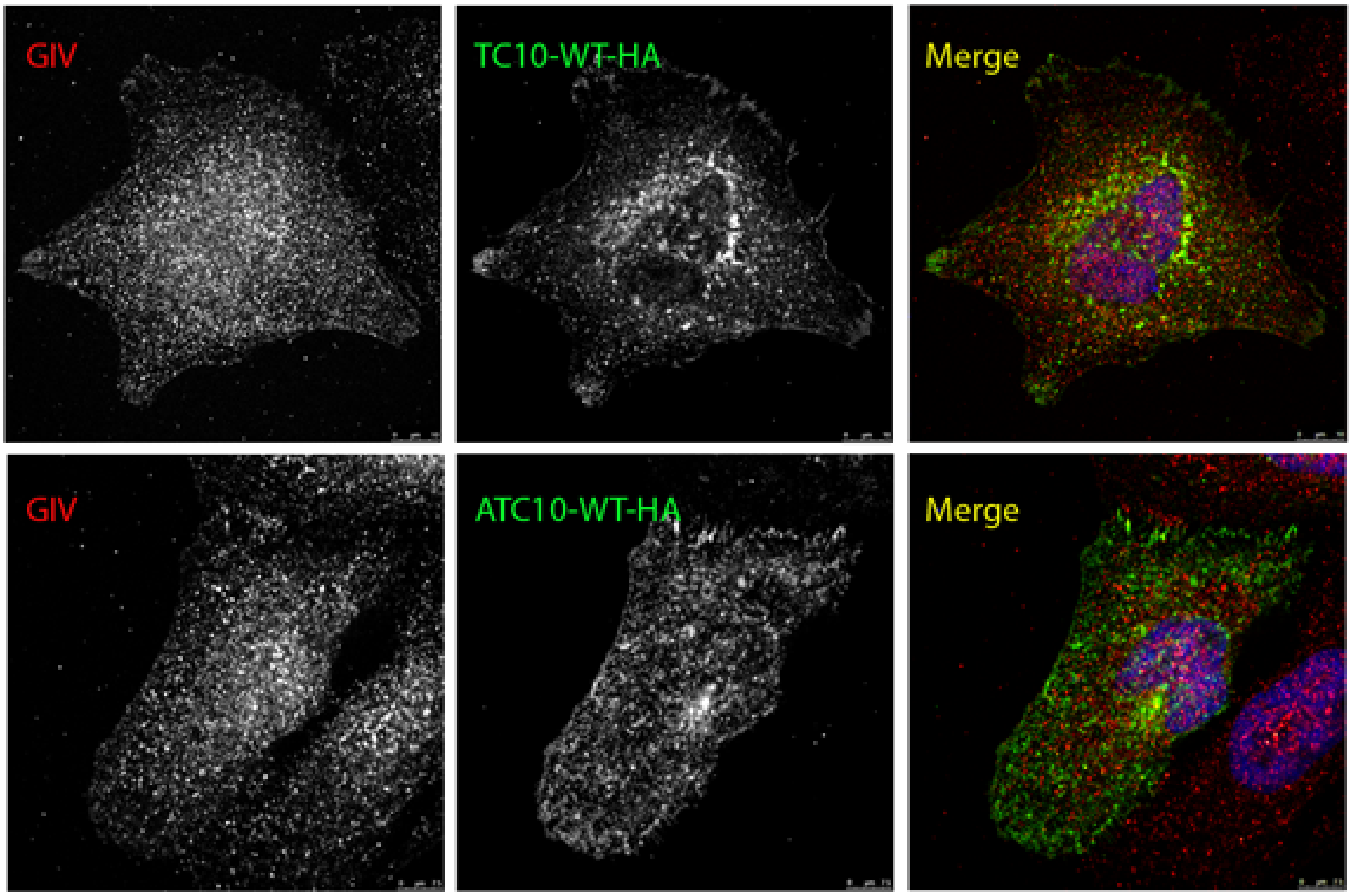


**Supplementary Figure 7 (related to Fig 6)**

**TC10-WT cannot recruit endogenous GIV to the PM sites.**

MDA MB231 cells exogenously expressing TC10-WT-HA (green) were fixed and stained for endogenous GIV (red). Representative images of transfected cells are shown. Red and green panels are shown in grey scale.

### SUPPLEMENTARY MOVIES AND LEGENDS

**Supplementary Movie 1 [Link]**

Time-lapse TIRF images of a Control (shControl) MDA-MB231 cell transiently expressing MT1-MMP-pHluorin shows real-time exocytic events. Images were acquired over 10 minutes. The movie was acqured at exposure time of 100 ms and with 1 pixel corresponding to 160 nm.

**Supplementary Movie 2 [Link]**

Time-lapse TIRF images of a GIV-depleted (by shRNA) MDA-MB231 cell transiently expressing MT1-MMP-pHluorin shows real-time exocytic events. Images were acquired over 10 minutes. The movie was acqured at exposure time of 100 ms and with 1 pixel corresponding to 160 nm.

**Supplementary Movie 3 [Link]**

Time-lapse TIRF images of a GIV-depleted MDA-MB231 cell stably expressing GIV-WT-FLAG and transiently expressing MT1-MMP-pHluorin shows real-time exocytic events. Images were acquired over 10 minutes. The movie was acqured at exposure time of 100 ms and with 1 pixel corresponding to 160 nm.

**Supplementary Movie 4 [Link]**

Time-lapse TIRF images of a GIV-depleted MDA-MB231 cell stably expressing a Exo70 binding-deficient GIV-AAxA-FLAG mutant and transiently expressing MT1-MMP-pHluorin shows real-time exocytic events. Images were acquired over 10 minutes. The movie was acqured at exposure time of 100 ms and with 1 pixel corresponding to 160 nm.

**Supplementary Movie 5 [Link]**

Time-lapse TIRF images of a GIV-depleted MDA-MB231 cell stably expressing another Exo70 binding-deficient GIV-F1742A-FLAG mutant MT1-MMP-pHluorin shows real-time exocytic events. Images were acquired over 10 minutes. The movie was acqured at exposure time of 100 ms and with 1 pixel corresponding to 160 nm.
